# Pattern-dependent low-intensity repetitive magnetic stimulation enhances spinal cord repair through modulation of neuroinflammation

**DOI:** 10.64898/2026.09.09.750117

**Authors:** Pauline Neveu, Alexandre Du, Fannie Semprez, Laurine Moncomble, Aurélien Fauquier, Camille Salat, Abigail Mc Gowan, Emma Darracq Mousli, Clémence Raimond, Rachel Sherrard, Fatiha Nothias, Nicolas Guérout

**Affiliations:** Saints-Pères Paris Institute for the Neurosciences, Université Paris-Cité, CNRS UMR 8003, Paris, France; Neural Adaptation and Repair, Development Adaptations and Aging, CNRS UMR 8263, INSERM U1345, IBPS, Sorbonne University, Paris, France

**Keywords:** spinal cord injury, repetitive magnetic stimulation, low intensity, neuroinflammation, non-invasive neuromulation

## Abstract

Spinal cord injuries (SCI) are traumatic lesions of the spinal cord that commonly result from physical trauma. They can lead to severe and permanent impairments in motor, sensory, and/or autonomic functions, often resulting in paraplegia or tetraplegia. Despite advances in this field, effective restorative treatments for SCI remain limited.

Over the past few years, several therapeutic strategies have been investigated to promote functional recovery after SCI, including neuromodulation approaches. Repetitive magnetic stimulation has emerged as a promising non-invasive strategy, with encouraging results when the stimulation is applied directly over the spinal cord. However, most studies have focused on high-intensity stimulation, while the key parameters underlying its therapeutic efficacy remain poorly understood. In this study, we investigated the effects of low-intensity repetitive trans-spinal magnetic stimulation (LI-rTSMS) following SCI in mice, using BMS, histological and RNA sequencing approaches. We aimed to determine the optimal protocol by comparing three stimulation patterns (10 Hz, iTBS and BHFS) and two coil sizes.

Our results show that LI-rTSMS modulates the injured spinal cord in a pattern- and coil size-dependent manner, with distinct effects on fibrotic, neural, and inflammatory responses. Notably, the BHFS pattern produced the most marked tissue-remodeling effects, reducing fibrosis, reactive astrogliosis, and phagocytosis of myelin debris. This reduction in phagocytosis was also observed with a smaller coil, supporting an effect of LI-rTSMS on this process. We also identified distinct inflammatory signatures depending on the stimulation pattern and time after injury. Transcriptomic analyses revealed a common early inflammatory response to LI-rTSMS, whereas each stimulation pattern subsequently led to distinct molecular signatures after 2 weeks stimualtion. In addition, ependymal cells are known to retain endogenous regenerative potential following SCI. We found that LI-rTSMS increased ependymal cell proliferation and migration toward the lesion site, without detectable changes in their differentiation.

Finally, we assessed c-Fos expression as a marker of early transcriptional changes in neurons. LI-rTSMS did not result in significant changes, suggesting that the observed effects were unlikely to be mediated by early, widespread neuronal activation.

Altogether, this study provides mechanistic insights into the pattern-dependent effects of LI-rTSMS and highlights its potential role for future clinical translation.

## Introduction

Spinal cord injury (SCI) is a devastating neurological condition that profoundly affects patients’ quality of life. According to the World Health Organization, approximately 15.4 million people worldwide are living with a SCI [1]. SCI can result from either traumatic – commonly arises from road traffic accidents, falls and violence – or non-traumatic etiologies which include inflammatory, ischemic and neoplastic causes [2,3]. Traumatic SCI remains the most common form reported worldwide and constitutes the vast majority of experimental models used to study spinal cord repair in rodents.

SCI frequently leads to permanent motor, sensory and autonomic deficits below the level of injury. Its pathophysiology is characterized by a primary injury, corresponding to the initial mechanical trauma, followed by a secondary phase characterized by spinal cord edema, ischemia, blood-spinal cord barrier disruption, axonal degeneration, neuronal apoptosis and excitotoxicity [4,5]. SCI is also associated with a robust inflammatory response involving microglia, the resident immune cells of the central nervous system, as well as infiltrating peripheral innate and adaptive immune cells including neutrophils, lymphocytes, and blood-derived monocytes that differentiate into macrophages sharing overlapping functions and phenotypes with microglia. Together, these cells orchestrate the inflammatory response, with microglia/macrophages (Mi/MΦ) predominantly adopting a pro-inflammatory phenotype during the acute phase following SCI [6,7].

As the lesion evolves, perivascular fibroblasts infiltrate the injury site and contribute to the formation of the fibrotic scar, replacing necrotic tissue and creating a microenvironment that is poorly permissive for axonal regeneration[8]. Recent lineage-tracing and transcriptomic studies have revealed that the fibrotic scar is heterogeneous, mainly composed of perivascular and meningeal fibroblasts that occupy distinct spatial niches and exert complementary functions during tissue remodeling [9]. In parallel, reactive astrocytes proliferate and contribute to the formation of the glial scar surrounding the lesion. This response initially protects the tissue by limiting the spread of inflammation and preserving the adjacent healthy tissue. However, the role of reactive astrocytes is highly context-dependent and remains debated. Anderson *et al.* demonstrated that their genetic ablation impaired axonal regrowth and worsened tissue repair following SCI [10]. Conversely, reactive astrocytes can also contribute to a non-permissive lesion environment through the secretion of inhibitory extracellular matrix molecules, including chondroitin sulfate proteoglycans. More recent evidence further suggests that astrocytes can promote tissue repair through the release of extracellular vesicles carrying neuroprotective factors, whereas distinct reactive astrocytes may exacerbate inflammation depending on their activation state [11].

The management of SCI represents a major economic and societal challenge. Current clinical care relies primarily on early surgical decompression to stabilize the vertebral column and reduce secondary damage, followed by intensive rehabilitation [12]. Although therapeutic approaches such as riluzole and methylprednisolone [13], cell transplantation [14,15] or tissue-engineering based on biomaterial implantation [16] have demonstrated tissue preservation and functional improvements in animal models, no treatment currently enables complete recovery of lost functions in humans. However, the spinal cord retains a limited endogenous capacity through ependymal cells lining the central canal. Following SCI, these cells become activated, proliferate, migrate toward the lesion site, and differentiate mainly into astrocytic, and to a less extent, oligodendrocytic lineages, contributing to tissue remodeling and scar formation [17,18]. This endogenous stem cell response represents a potential therapeutic target to enhance spinal cord repair.

In recent years, several neuromodulatory approaches have emerged as promising strategies to modulate the central nervous system activity, promote functional recovery and enhance endogenous repair mechanisms, including axonal plasticity and modulation of neuroinflammation [19–21]. Among these approaches, electrical stimulation has been extensively investigated to treat SCI, using both invasive and non-invasive strategies. In patients with chronic SCI, epidural electrical stimulation has yielded promising clinical outcomes, including restoration of voluntary locomotion and, more recently, brain-controlled walking through its integration with brain-computer interfaces [22,23]. Another major neuromodulatory approach is repetitive transcranial magnetic stimulation (rTMS), a non-invasive technique based on Faraday’s principle of electromagnetic induction. A stimulator generates rapidly changing electrical currents within a coil, producing a magnetic field that induces secondary electrical currents in the underlying neural tissue. Applied over the brain, rTMS has therapeutic efficacy in several neurological disorders, such as stroke and Parkinson’s disease [24], as well as psychiatric disorders. More recently, accelerated intermittent theta burst stimulation (iTBS), a standardized pattern of rTMS, has reduced depressive-like behaviors in mice [25].

These advances have prompted the development of magnetic stimulation protocols directly targeting the spinal cord. Repetitive trans-spinal magnetic stimulation (rTSMS) became a promising non-invasive therapeutic strategy for spinal cord disorders. In a mouse model of focal demyelination, we previously demonstrated that rTSMS decreased neuroinflammation and demyelination in a sex-dependent manner [26]. Similarly, Mi/MΦ rTSMS promotes functional recovery following SCI in mouse transection models as well as in rat transection and contusion models, reducing motor deficits while modulating the fibroglial scar and inflammation [27–29]. rTSMS also regulates the response of Mi/MΦ, enhancing their phagocytic activity, promoting myelin debris clearance and axonal repair [30,31]. Moreover, we previously reported that rTSMS improves the self-renewal, migration and differentiation of ependymal cells following SCI, suggesting that it may promote spinal cord repair through activation of endogenous regenerative mechanisms [27].

However, conventional rTSMS is generally delivered at high-intensities, resulting in relatively diffuse stimulation that limits spatial specificity and complicates translation across experimental models. Low-intensity rTMS (LI-rTMS) has recently emerged as a promising alternative, enabling more focal magnetic stimulation. Dufor *et al.* demonstrate that LI-rTMS applied to the cerebellum induces dendritic spine reorganization and reinnervation of Purkinje cells, they also identify cryptochrome as a putative magnetoreceptor mediating the biological effects of magnetic stimulation [32]. LI-rTMS reduces the expression of genes associated with inflammation and calcium signaling in cultured mouse cortical astrocytes [33], and very low-intensity static magnetic stimulation improves locomotor recovery, increases tissue sparing, and reduces calcium channel expression in a rat model of SCI [34].

Because neuronal and glial cells can exhibit frequency-dependent responses to electromagnetic stimulation, stimulation patterns may engage distinct molecular pathways, modulate neuronal plasticity, and recruit different cellular populations. Thus, the biological effects of LI-rTMS may critically depend on the stimulation parameters. Complex patterns may therefore provide an opportunity to fine-tune the biological response to low-intensity stimulation. For instance, Sherrard and colleagues developed a biomimetic high-frequency stimulation (BHFS) pattern designed to mimic endogenous electrical activity generated around peripheral nerves during exercise [35,36]. Although conventional rTSMS has shown promising effects on tissue repair and functional recovery following SCI, the effects of low-intensity magnetic fields remain poorly characterized, and whether complex patterns can differentially modulate tissue repair are largely unknown.

To address this question, we developed a low-intensity rTSMS (LI-rTSMS) device designed for focal stimulation of the injured spinal cord. Using three distinct stimulation patterns (10 Hz, iTBS and BHFS) and two coil sizes, we investigated their effects on locomotor recovery, tissue remodeling, and inflammatory responses following SCI. We further characterized the transcriptional responses associated with each stimulation pattern and investigated whether LI-rTSMS could modulate the endogenous regenerative potential of the injured spinal cord by influencing the response of ependymal cells.

## Material and methods

### Animal care and use statement

All experimental procedures complied with the European Community guidelines for the care and use of laboratory animals (86/609/EEC; Official Journal of the European Communities No. L358, Decembre 18, 1986), The French Decree No. 87/848 of October 19, 1987, and were approved by the local Ethics Committee (#202307121857640). All protocols were designed to minimize animal use and suffering.

Experiments were performed on adult male and female C57BL/6J mice, or on tamoxifen inducible hFoxJ1-CreERT2::eYFP transgenic mice generated by crossing heterozygous hFoxJ1-CreERT2 mice with Rosa26-eYFP reporter mice. To induce Cre recombination, tamoxifen (75 mg/kg; 20 mg/mL, Sigma-Aldrich, T5648-1G; approximately 90 µL for a 25 g mouse) dissolved in corn oil (Fisher, 405435000) was administered by four consecutive daily subcutaneous injections. SCI was performed 10 days after the last tamoxifen injection to ensure sufficient eYFP expression in FOXJ1 expressing cells and to avoid unspecific recombination. Mice (7–12 weeks of age; *n* total = 127) were group-housed (five animals per cage) in a conventional rodent facility (BiomedTech Facilities, Université Paris Cité) under controlled environmental conditions.

### Spinal cord injury

All mice underwent a SCI, except for a separate cohort used for the c-Fos experiment which underwent laminectomy only and control animals in the ependymal cell experiment. Briefly, mice were deeply anesthetized with isoflurane (5 % in oxygen at 3 L/min) and premedicated by subcutaneous injection of buprenorphine (0.03 mg/mL in 0.9 % NaCl, 3 µL/g body weight). Anesthesia was maintained using a mask (1.5–2 % isoflurane in oxygen at 1 L/min). Ophthalmic gel was applied to prevent corneal drying, and the dorsal surface was shaved and disinfected with povidone-iodine solution.

Following skin incision and muscle dissection, a laminectomy was performed to expose the spinal cord at the T11–T13 vertebral level (corresponding to the T9 spinal segment). The spinal cord of C57BL/6J mice was completely transected with a 25-gauge needle, and for the hFoxJ1-CreERT2::eYFP mice, a dorsal hemisection was carried out using a 30-gauge needle to preserve the integrity of the central canal as described previously [37]. Muscles and skin were sutured separately, and the surgical site was disinfected. Animals received a subcutaneous injection of sterile saline to compensate for fluid loss during surgery, and moist food pellets were placed on the cage floor to facilitate feeding during recovery. Buprenorphine was administered again at the end of the day, and the following day if signs of discomfort or pain.

### Therapy by low-intensity repetitive trans-spinal magnetic stimulation

LI-rTSMS was delivered using custom-designed circular coils specifically adapted for focal stimulation in rodents. Two coils were used:

- **Small coil:**400 turns of 35 AWG copper wire (external diameter: 0.15 mm; copper core area: 0.017 mm²). The coil core consisted of an inner cylinder with a diameter of 5.5 mm, a base diameter of 8 mm. Resistance: ∼ 9.6 Ω; inductance ∼ 2.3 mH.

- **Medium coil**: 800 turns of 31 AWG copper wire (external diameter: 0.22 mm), with a 2 mm ferrite core. The coil had an inner diameter of 6 mm, an outer diameter of 16 mm. Resistance: ∼ 9 Ω; inductance: ∼ 15.3 mH.

Both coils were connected to a custom-built stimulator delivering LI-rTSMS (peak magnetic field of 9 mT), powered by 9 V or 12 V adaptors, respectively to the size. The coil was placed over the lesion site on the back of the isoflurane anesthetized mouse to ensure focal and reproducible stimulation, temperature did not exceed 37 °C at skin contact. LI-rTSMS started 1 day post-injury (dpi) and was applied once daily (10 min/session) until tissue collection. Depending on the experimental paradigm, mice received 3, 6, or 14 consecutive stimulation sessions and were euthanized the day after the final stimulation session, except for the c-Fos experiment. SCI mice were randomly assigned to receive LI- rTSMS using either the small, the medium coil or sham stimulation under identical anesthesia conditions. Group allocation was balanced according to the BMS score at 1 dpi. Three patterns of LI- rTSMS were tested (Fig. 1A):

**Fig. 1.**
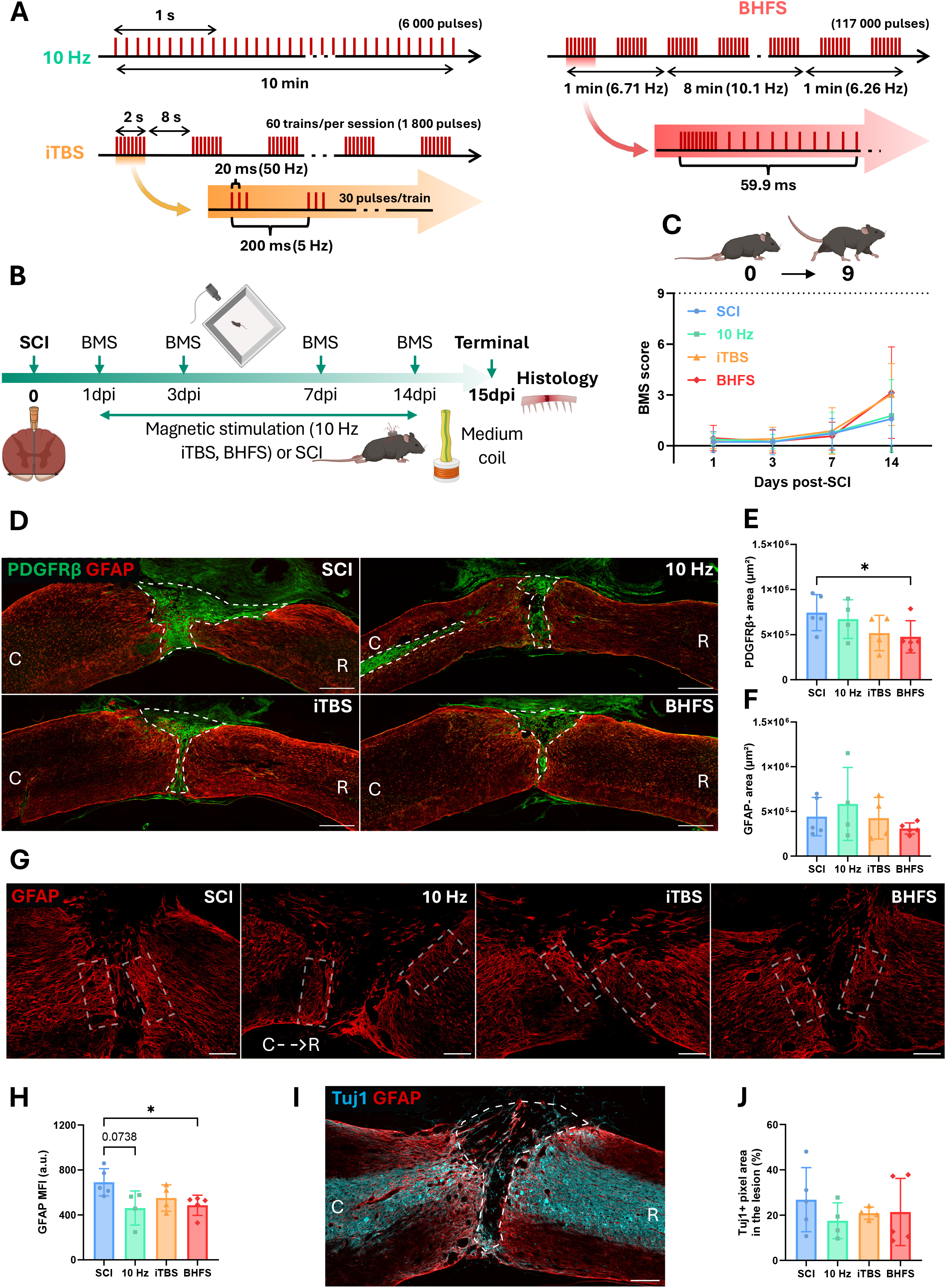
LI-rTSMS does not alter locomotor recovery, but the BHFS pattern reduces fibrocyte scar formation and astrocyte reactivity. **A.** Schematic representation of the three LI-rTSMS stimulation patterns used in this study for a 10 min session: **10 Hz** (top left), **iTBS** (bottom left), **BHFS** (top right). **B.** Experimental design: all mice underwent a SCI on day 0, then 3 groups received LI-rTSMS treatment from 1 to 14 dpi, using either the 10 Hz, iTBS, or BHFS stimulation pattern, while SCI animals without stimulation served as controls. In parallel, locomotor behaviors were recorded for all mice at 1, 3, 7 and 14 dpi, and animals were perfused at 15 dpi for histological analyses. **C.** Assessment of locomotor recovery over time using the BMS score. The white dotted line represents the baseline. **D–F.** Analysis of the effects of LI-rTSMS on fibroglial scar formation. **D.** Representative epifluorescence images (10× magnification) of sagittal spinal cord sections stained with anti-PDGFRβ (green) and anti-GFAP (red) antibodies, from top left to bottom right: SCI, 10 Hz, iTBS, and BHFS. The white dotted lines surround the lesion core. Scale bar: 500 µm. Quantifications of **E.** PDGFRβ-positive area (µm²) and **F.** GFAP-negative area (µm²). **G–H**. Analysis of the effects of LI-rTSMS on astrocytes reactivity at the scar borders. **G.** Representative confocal images (20× magnification) of sagittal spinal cord sections stained with anti-GFAP (red) antibody, from left to right: SCI, 10 Hz, iTBS, and BHFS. The white dotted rectangles surround the analysed scar borders. Scale bar: 200 µm. **H.** Quantification of GFAP mean fluorescence intensity (MFI) at the scar borders. **I–J.** Analysis of the effects of LI-rTSMS on axonal survival/regrowth within the lesion. **I.** Illustrative epifluorescence image (20× magnification) of a sagittal spinal cord section stained with anti-Tuj1 (cyan) and anti-GFAP (red) antibodies from a BHFS-treated mouse. The white dotted line surrounds the GFAP-negative lesion area. Scale bar: 200 µm. **J.** Quantification of Tuj1-positive pixel area within the GFAP-negative lesion area (%). Orientation: caudal (C) on the left and rostral (R) on the right in all sagittal sections. Statistical analyses compared the SCI group with each stimulation group using a Mixed-effects analysis with Šidák correction for multiple comparisons (**C**: SCI *n* = 11; 10 Hz *n* = 10; iTBS *n* = 10; BHFS *n* = 11), and the Kruskal-Wallis test with Dunn’s multiple-comparisons test (**E, F, H, J**: SCI *n* = 5; 10 Hz *n* = 4; iTBS *n* = 4; BHFS *n* = 5). Trend *p* value is indicated on the graph (**H**). (* = *p* ≤ 0.05). Created with BioRender.com.

- **10 Hz** (6 000 pulses): continuous stimulation at 10 Hz. 10 Hz is frequently used in human rTMS

- **iTBS** (1 800 pulses): bursts of 3 pulses at 50 Hz delivered at 5 Hz for 2 s, followed by 8 s of rest, repeated throughout the 10 min session.

- **BHFS** (117 000 pulses): 62.6 ms trains of 20 pulses repeated at 9.75 Hz. This complex pattern (patent PCT/AU2007/000454, Global Energy Medicine) was adapted to deliver LI-rTSMS in three phases: 1 min warm-up (6.71 Hz), 8 min treatment (10.1 Hz), and 1 min cooldown (6.26 Hz) [36].

### Open field test and locomotion recovery

To assess functional recovery, mice were filmed from above in an open field chamber for 3 min, one week before surgery (baseline), and at 1, 3, 7, and 14 days post-SCI. Videos were scored by one or two experimenters blinded to the experimental conditions using the Basso mouse scale, ranging from 0 (complete hindlimb paralysis) to 9 (normal locomotion) [38].

### Assessment of neuronal activity following LI-rTSMS

To evaluate acute neuronal activation induced by LI-rTSMS, a separate cohort of mice underwent laminectomy without SCI (*n* = 5 per group). One day later, animals received a single 10 min session of BHFS or Sham stimulation delivered using either the small or medium coil under isoflurane anesthesia. They remained anesthetized for 1 h following the onset of stimulation to minimize c-Fos induction associated with spontaneous behaviors such as locomotion or grooming. Mice were transcardially perfused, and cervical, thoracic and lumbar spinal cord segments were collected for immunohistochemistry as described below.

### Tissue preparation and immunohistochemistry

For histological experiments, mice received an intraperitoneal injection of a ketamine/xylazine mixture (ketamine: 0.75 µL/g; xylazine: 0.5 µL/g; diluted in 0.9 % NaCl; 100 µL/25 g body weight). They were transcardially perfused with 1X phosphate-buffered saline (PBS) containing EDTA (final concentration at 0.005 M), followed by 4 % paraformaldehyde (PFA; Diapath France, Antigenfix, P0014/U). Spinal cords were dissected, post-fixed overnight in 4 % PFA at 4 °C and cryoprotected in 30 % sucrose (Euromedex, 200-301-B) for 48 h. Samples were embedded in Tissue-Tek OCT compound (Epedria, Neg-50, 6502) in cryomolds and rapidly frozen in isopentane cooled to −30 °C to −50 °C. Tissues were stored at −80 °C until sectioning. Sagittal (20 µm) and transverse (20 or 30 µm) cryosections were cut using a cryostat and mounted onto SuperFrost Plus microscope slides (Thermo Fisher, J1800AMNZ), then stored at −20 °C until use for immunohistochemistry.

For staining, slides were thawed at room temperature and tissue sections were circled using a hydrophobic barrier pen (PAP pen, ClearLine, 490001), then placed in a humidified chamber. After rehydration in PBS, sections were incubated in 0.1 M glycine (Euromedex, 26-128-6405-C) for 20 min to reduce background, followed by a blocking step in PBS containing 0.5 % Triton X-100 (Euromedex, 2000-B) and 1X commercial blocking solution (Abcam, ab126587) for 1 h at room temperature. Primary antibodies were diluted in blocking solution and incubated overnight at 4 °C. After washing in PBS containing Tween-20 (0.05 %; Euromedex, 2001-C), sections were incubated with secondary antibodies (1:500) in a blocking solution for 1 h at room temperature. Nuclei were counterstained with **DAPI** (Invitrogen, D3571; 1:5000). Slides were cover-slipped using Mowiol mounting medium.

The following primary antibodies were used: **GFAP** (mouse monoclonal IgG1 conjugated to Cy3, Sigma-Aldrich, C9205; 1:500). **GFAP** (chicken polyclonal IgY, Abcam, ab4674; 1:500). **PDGFRβ** (rabbit monoclonal IgG, Abcam, ab32570; 1:500). **Iba1** (rabbit monoclonal IgG1, Synaptic Systems, 234008; 1:1000). **MBP** (rat monoclonal IgG2a, Sigma-Aldrich, MAB386; 1:500). **Tuj1** (rabbit polyclonal IgG, BioLegend, 802001; 1:2000). **NeuN** (mouse A60 monoclonal conjugated to Alexa Fluor 488, Sigma-Aldrich, MAB377X; 1:200) or **NeuN** (rabbit polyclonal IgG, Abcam, ab104225; 1:500). **c-Fos** (guinea pig monoclonal IgG2, Synaptic Systems, 226 308; 1:500) or **c-Fos** (rabbit monoclonal IgG, Synaptic System, 226 008; 1:1000). **GFP** (chicken polyclonal IgY, Aves Lab, GFP-1020; 1:5000) or **GFP** (rabbit polyclonal IgG conjugated to Alexa Fluor 488, Jackson immunoresearch, 300-545-245; 1:1000). **SOX9** (goat polyclonal IgG, R&D, AF3075; 1:300). **SOX10** (goat polyclonal IgG, R&D, AF2864, 1:500). **Ki67** (rabbit polyclonal IgG, Abcam, ab15580-1001; 1:500). Secondary antibodies were used in donkey or goat, and conjugated to Alexa Fluor 488, 555, 633 or 647, or to Cyanine-3 (Invitrogen, Thermo fisher).

### Microscopy imaging and quantification

Microscopy images were acquired using an epifluorescence scanner (Zeiss AxioScan.Z1, ZEN 3.1 software) or confocal microscopes (Zeiss LSM 710 and 880, ZEN Black 2.1 software). Image analysis was performed using QuPath (v0.7.0) and Fiji (ImageJ) by experimenters blinded to the experimental groups. Spinal cord sections were imaged at 10× or 20× magnifications. Z-stack images were acquired on selected sections and processed as maximum intensity projections using Fiji. For each animal, two to five sections encompassing the lesion epicenter were examined.

The section exhibiting the largest GFAP-negative area (astrocytic defect) and PDGFRβ-positive area (fibrotic scar) was selected for quantitative analyses. To assess glial scar mean fluorescence intensity (MFI), two rectangular regions of interest (ROI; 0.5 x 0.2 mm) were positioned in the rostral and caudal parts of the GFAP-positive scar surrounding the lesion. Tuj1 immunoreactivity was quantified using a machine learning–based pixel classifier within the GFAP-negative lesion area.

The same strategy was applied for Iba1, MBP and CD68 stainings by selecting the section exhibiting the largest areas of Iba1-positive (Iba1⁺) cells, CD68-positive (CD68⁺) cells, and MBP-negative area. Myelin debris was quantified as MBP-positive pixels within the lesion. To assess myelin debris clearance by Mi/MΦ, Iba1⁺ or CD68⁺ cells were detected using DAPI-based nuclear segmentation combined with a machine learning classifier. Cells containing MBP-positive signals were identified and expressed as the percentage of total Iba1⁺ or CD68⁺ cells [31]. Iba1 and CD68 MFI were quantified within the spinal cord in a single ROI, placed consistently around the lesion core across animals.

For the neuronal activity experiment, NeuN-positive (NeuN⁺) and c-Fos-positive (c-Fos⁺) cells were detected using DAPI-based nuclear machine learning. Co-labeled NeuN⁺c-Fos⁺ cells were expressed as the percentage of total NeuN⁺ neurons within the same section. Quantifications were performed across the whole spinal cord, subdivided by regions (cervical, thoracic, and lumbar), as well as separately in the dorsal and ventral horns.

The maximum migration distance of YFP-positive (YFP⁺) cells was measured as the distance between the central canal and the farthest YFP⁺ cell located within the glial scar. Migrating YFP⁺ area was calculated as the marker area within the glial scar. Nuclei were detected and YFP⁺ cells were counted in the spinal cord. To assess ependymal cell proliferation, three regions encompassing the lesion (rostral, lesion, and caudal) were analyzed. At least two sections per region were quantified by counting the total number of YFP⁺ cells lining the central canal and expressing Ki67. SOX9 immunofluorescence intensity was quantified from confocal images by measuring the fluorescence signal within a manually delineated ROI encompassing the central canal, after subtraction of the background signal. Ependymal cell differentiation was assessed using SOX9 and SOX10 as markers of astrocytic and oligodendrocytic lineages differentiation, respectively. SOX9-positive and SOX10- positive nuclei were identified within YFP⁺ regions using a machine learning–based nuclear classifier, with DAPI staining used for nuclear segmentation.

### Transcriptomic analysis

#### RNA sequencing and bioinformatic analysis

The LI-rTSMS medium coil was used for transcriptomic analysis. Six mice per group (SCI, 10 Hz, iTBS, BHFS) were sacrificed 24 h after the last session of stimulation, at either 4 or 15 days post-SCI. Animals in the 15 dpi group also performed the BMS and were included in Figure 1. All the mice were transcardially perfused with sterile 1X PBS under semi-sterile conditions by two experimenters. Spinal cord segments spanning the lesion site, 5 mm rostral and 5 mm caudal to the injury, were dissected, snap-frozen in liquid nitrogen, and stored at −80 °C until processing. Total RNA was extracted and quality-controlled on the BGI Tech Solutions platform (Hong Kong, China) using the RNA Integrity Number (RIN). Most samples showed high RNA quality, with RIN values ranging from 7.0 to 9.9, while two samples exhibited lower RIN values (5.1 and 5.3).

Poly(A)+ mRNA was enriched from total RNA using oligo(dT)-coupled magnetic beads and fragmented prior to cDNA synthesis using random hexamer primers. Double-stranded cDNA libraries were generated following end repair, adapter ligation, and PCR amplification. Library quality was assessed before sequencing, which was performed on the DNBSEQ™ platform (BGI Genomics) using paired-end 150 bp reads. Raw sequencing reads were filtered to remove adapter sequences, low-quality reads, and reads containing excessive numbers of unknown bases. Clean reads were aligned to the *Mus musculus* reference genome (GRCm39) using HISAT2 and Bowtie2 [39,40]. Alignment quality was evaluated based on mapping statistics and read distribution across the reference genome.

Gene expression count matrices were generated using the Dr. Tom platform (BGI Genomics) and imported into the R Studio statistical environment for downstream analyses. Principal component analysis, sample correlation and differential gene expression analyses were performed on normalized gene expression data. Differentially Expressed Genes (DEGs) were identified using the DEGSeq package in R Studio [41]. Genes with a false discovery rate < 0.05 were considered significantly differentially expressed, with an absolute log2 fold change > 1. DEGs were subsequently classified as upregulated or downregulated according to the direction of the observed expression changes.

To investigate the biological significance of the DEGs, Gene Ontology (GO) enrichment analyses were performed. GO terms were categorized into biological process, molecular function, and cellular component. A statistical enrichment test was applied to determine significantly associated terms, with correction for multiple comparisons. Because the primary objective of this study was to identify biological pathways associated with SCI and the effects of neuromodulation, the biological process category was prioritized for interpretation. GO enrichment analyses were visualized using the Cytoscape software (v3.10.3) with ClueGO (v2.5.10) and CluePedia (v1.5.10) plugins. Functionally related GO terms were grouped into clusters and displayed as interaction networks (Figs. 3–4). Within each cluster, the most significantly enriched term was designated as the leading term. Node size was proportional to enrichment significance, whereas edges represented functional relationships between pathways. GO networks were generated using the top 400 DEGs from each comparison, corresponding approximately to genes displaying an absolute fold change greater than 50 %. Detailed analysis parameters are provided in Supplementary Table 1, and the GO terms used for GO analysis results, including associated genes, are provided in the accompanying Supplementary Table S2. Overlaps between GO terms identified across the different conditions are shown as Venn diagrams (Figs. S7–S8).

To obtain a broader overview of transcriptional changes, Gene Set Enrichment Analysis (GSEA) was performed using ranked gene expression data from all detected genes [42]. Gene sets from the Molecular Signatures Database (MSigDB), including GO biological process and pathway-related collections, were analyzed to identify significantly enriched biological pathways. GSEA results were subsequently imported into Cytoscape using the EnrichmentMap (v3.5.0) plugin to generate pathway interaction networks based on gene overlap and functional similarity. To facilitate network interpretation, the AutoAnnotate (v1.5.2) plugin automatically groups related pathways into thematic clusters according to semantic similarity (Figs. S1−S6). DEG, GSEA, and Cytoscape data are provided in the corresponding Excel file for each comparison (Tables. S3−S5).

### Statistics

All statistical analyses were performed using GraphPad Prism (v11.0.2). Data normality was assessed with the Shapiro–Wilk test. Comparisons between two groups were performed using an unpaired Student’s *t*-test for normally distributed data or the Mann–Whitney test for non-normally distributed data. Comparisons among more than two groups were performed using one-way ANOVA followed by Dunnett’s multiple comparisons test for normally distributed data, or the Kruskal–Wallis test followed by Dunn’s multiple comparisons test for non-normally distributed data. Depending on the experimental design, two-way ANOVA or a Mixed-effects model were used, followed by Dunnett’s multiple comparisons test when appropriate. Spearman’s rank correlation and simple linear regression were performed as appropriate for correlation analyses. Data are presented as mean ± standard deviation (SD). A *p* value ≤ 0.05 was considered statistically significant.

## Results

### LI-rTSMS patterns differentially regulate fibroglial scar formation, Mi/MΦ response and myelin debris clearance

Mice underwent SCI and received daily LI-rTSMS treatment starting 1 dpi and continuing for 14 consecutive days. Magnetic stimulation was delivered for 10 min per day using a medium-sized coil (1.6 cm diameter), and we evaluated three distinct stimulation patterns: 10 Hz, iTBS and BHFS (Fig. 1A). Locomotor recovery was assessed at 1, 3, 7 and 14 dpi using the BMS, following a 3 min recording of spontaneous locomotion in an open field. At 15 dpi, animals were perfused and spinal cords were collected for histological analyses (Fig. 1B).

Locomotor assessment revealed that, at the early time points of 1, 3 and 7 dpi, all groups exhibited comparable BMS scores (notably a score of 0 at 1 and 3 dpi), reflecting both the severity of the lesion and the homogeneity of the injury across experimental groups (Fig. 1C). At 14 dpi, mice receiving iTBS or BHFS stimulation displayed a greater degree of locomotor recovery than SCI and 10 Hz-treated animals, although these differences did not reach statistical significance. In particular, 18 % of mice in the SCI group reached a BMS score ≥ 3 (corresponding to plantar placement, with or without weight support), compared to 30 % in the 10 Hz group, 50 % in the iTBS group, and 45 % in the BHFS group (*n* = 10–11 mice per group) (Fig. 1C).

Spinal cords were subsequently analyzed by immunohistochemistry to assess the fibroglial scar (Fig. 1D–H). These analyses demonstrated that only LI-rTSMS delivered using the BHFS pattern significantly reduced the PDGFRβ-positive fibrotic scar area within the lesion core compared with SCI controls (Fig. 1E). Concerning the astroglial component of the scar, the GFAP-negative area within the lesion core was quantified, no stimulation paradigm significantly affected this parameter compared with SCI animals (Fig. 1F). However, analysis of astrocyte reactivity at the lesion border, assessed by GFAP fluorescence intensity, revealed that both the 10 Hz and BHFS stimulation protocols significantly reduced astrogliosis (Fig. 1G–H).

These analyses were complemented by the evaluation of axonal survival/regrowth using Tuj1 immunolabeling. No significant differences were observed between stimulated groups and SCI controls for this parameter (Fig. 1I–J).

Several studies have investigated the effects of rTMS in neurodegenerative and traumatic disorders, and have reported anti-inflammatory responses [24]. It was recently demonstrated that rTMS directly modulates Mi/MΦ activity via the p62/Nrf2 and p38 MAPK pathways, and enhances myelin debris clearance at the lesion site after SCI [31,43]. We therefore investigated the effects of LI-rTSMS stimulation patterns on the inflammatory processes, myelin debris accumulation and clearance at 15 days post-SCI (Fig. 2). We first performed a spatial analysis of Iba1⁺ pixel area along the rostro-caudal axis, from 2.4 mm caudal to 2.55 mm rostral to the lesion epicenter, to map Mi/MΦ distribution throughout the injured spinal cord (Fig. 2A–C). This analysis revealed a progressive increase in Iba1⁺ pixel area toward the lesion epicenter in all experimental conditions. However, no marked differences were observed between stimulation paradigms (Fig. 2B–C). We subsequently assessed Mi/MΦ reactivity by quantifying Iba1 fluorescence intensity throughout the lesion area on confocal images. BHFS increased Iba1 fluorescence intensity compared with SCI controls (Fig. 2D, G).

**Fig. 2.**
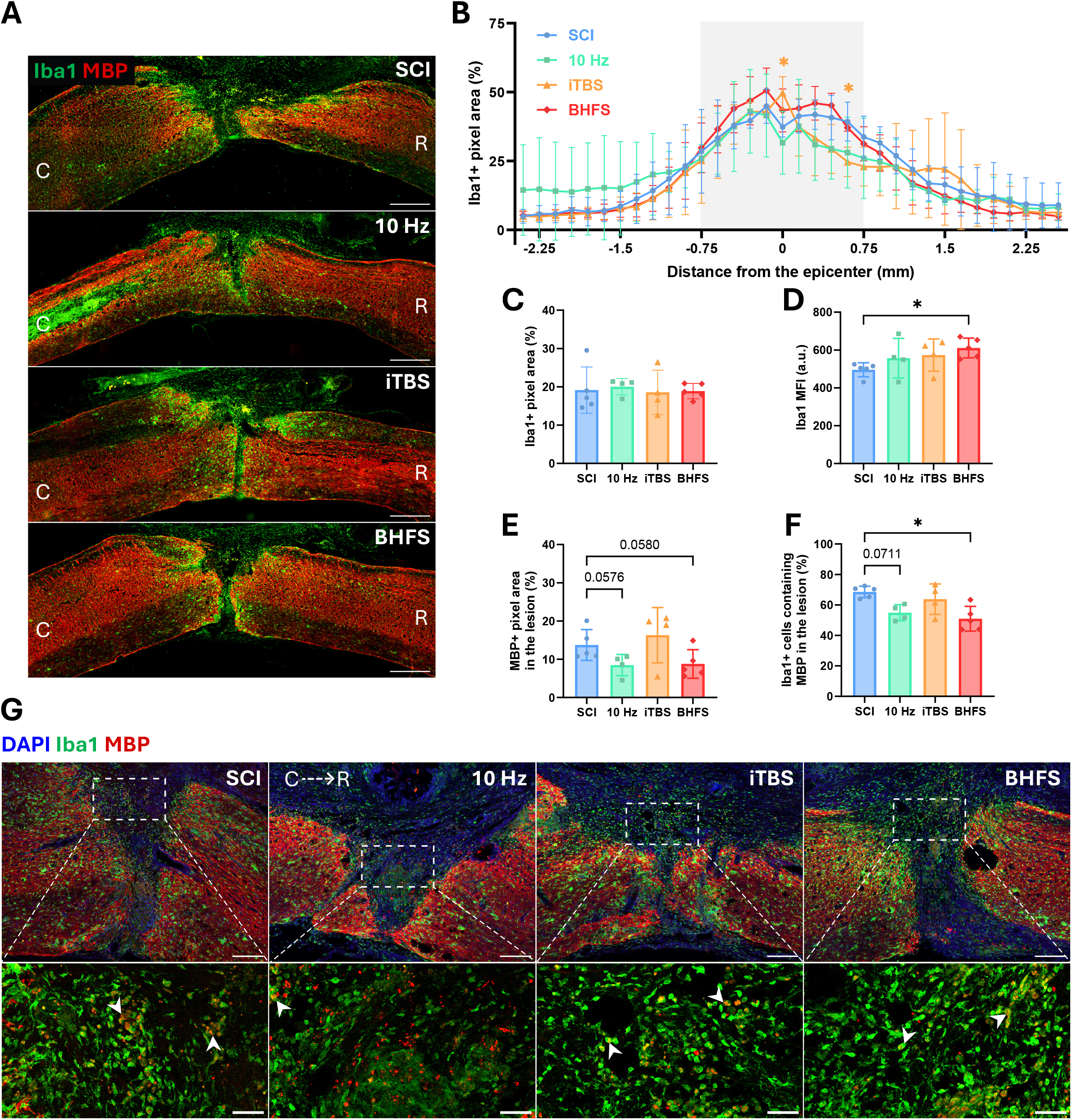
BHFS pattern enhances inflammatory reactivity and reduces their myelin debris clearance within the lesion. **A–G**. LI-rTSMS effects on inflammation and clearance of myelin debris at 15 days post-SCI. **A**. Representative epifluorescence images (10× magnification) of sagittal spinal cord sections stained with anti-Iba1 (green) and anti-MBP (red) antibodies, from top to bottom: SCI, 10 Hz, iTBS, and BHFS. Scale bar: 500 µm. **B**. Quantification of Iba1⁺ pixel area (%) from -2.4 mm to +2.55 mm around the lesion epicenter. The light grey-shaded area represents the lesion epicenter. **C**. Quantification of the average Iba1⁺ pixel area (%) across the entire spinal cord. **D**. Quantifications of Iba1 MFI. **E**. Quantifications of MBP⁺ pixel area within the MBP-negative lesion area (%), and **F**. the proportion of Iba1⁺ cells containing MBP⁺ debris inside the lesion (%). **G**. Representative confocal images (20× magnification) of sagittal spinal cord sections stained with DAPI (blue), anti-Iba1 (green) and anti- MBP (red) antibodies, from left to right: SCI, 10 Hz, iTBS, and BHFS. Scale bar: 200 µm. The white dotted rectangles indicate the regions zoomed in each panel below, with white arrows pointing to colabeled cells. Scale bar: 50 µm. Orientation: caudal (C) on the left and rostral (R) on the right in all sagittal sections. Total number of mice: SCI *n* = 5; 10 Hz *n* = 4; iTBS *n* = 4; BHFS *n* = 5. Statistical analyses compared the SCI group with each stimulation group using a Two-Way ANOVA with Dunnett’s correction for multiple comparisons (**B**), and the Kruskal-Wallis test with Dunn’s correction for multiple comparisons (**C–F**). In **B**, the asterisk is shown in the color corresponding to the stimulation group for which a significant difference was detected compared to the SCI group. Trend *p* values are indicated on the graphs (**E–F**). (**\*** = *p* ≤ 0.05).

Then, both BHFS and 10 Hz showed a trend toward a reduced myelin debris area within the MBP-negative lesion area compared with SCI controls (*p = 0.0580* and *p = 0.0576*, respectively; Fig. 2E). To further characterize the association between Mi/MΦ and myelin debris, we quantified the proportion of Iba1⁺ cells containing MBP within the lesion area. This parameter therefore reflects the proportion of Mi/MΦ containing detectable myelin debris. Interestingly, the percentage of phagocytic Iba1⁺ cells was significantly reduced in the BHFS-treated group and showed a tendency to decrease in the 10 Hz group (Fig. 2F).

Altogether, these results demonstrate that LI-rTSMS can modulate fibroglial scar formation, inflammatory response and myelin debris clearance following SCI. Importantly, these effects appear to be highly dependent on the stimulation pattern applied.

### Distinct transcriptional signatures emerge across stimulation patterns and post-injury stages

LI-rTSMS had limited effects on motor recovery but exerted more pronounced changes on spinal tissue (Figs. 1–2). We performed RNA sequencing (RNA-seq) and GO enrichment analyses to identify genes modulated by LI-rTSMS compared with the SCI group (Figs. 3–4) and then confirmed them through GSEA analyses of altered cellular processes (Figs. S1–S6). We identified cellular processes activated by LI-rTSMS after 3 daily stimulation sessions, as well as those sustained until the end of the 14 day treatment, which correspond to 4 days (Fig. 3) and 15 days post-SCI (Fig. 4), respectively. Because LI- rTSMS therapy altered the expression of numerous genes, we focused our analyses on the top 400 DEGs (Table. S2).

**Fig. 3.**
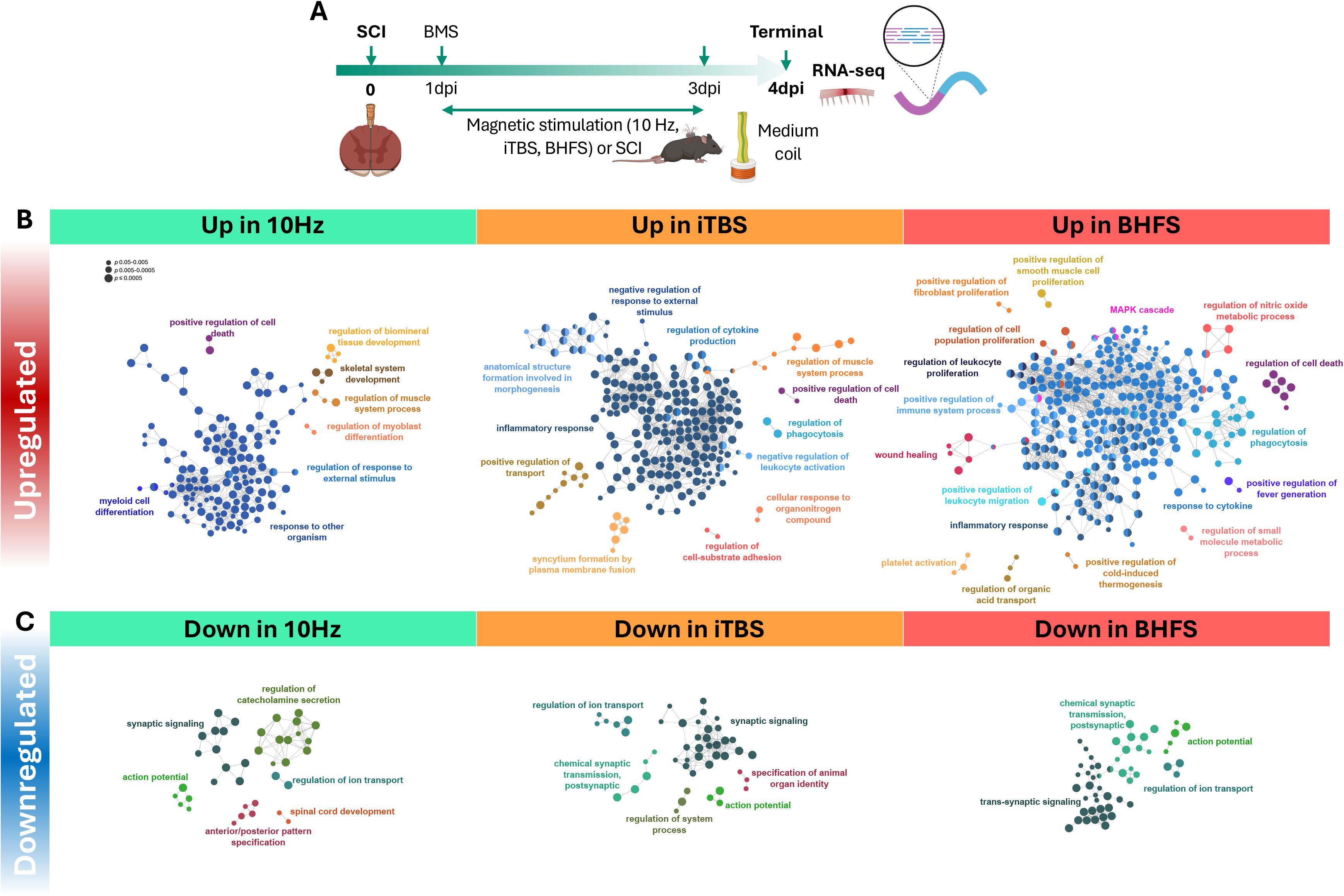
Transcriptomic sequencing revealed a similar profile between therapies with a reduction of inflammation at 4 dpi. **A**. Experimental design: all mice underwent a SCI on day 0, then 3 groups received LI-rTSMS treatment from 1 to 3 dpi using either the 10 Hz, iTBS, or BHFS stimulation pattern, while SCI animals without stimulation served as controls. The BMS score was assessed at 1 dpi to confirm injury severity consistency across groups. All animals were sacrificed at 4 dpi for RNA-seq analyses. **B–C**. ClueGO was used to identify significantly enriched Gene Ontology (GO) biological processes among upregulated (**B**) and downregulated (**C**) genes in each experimental condition compared to the SCI control. Bubble charts represent enriched GO terms (from left to right: 10 Hz, iTBS, and BHFS), where node size is proportional to the statistical significance of enrichment (*p* values range from ≤ 0.05 to ≤ 0.0005, as indicated by the scale). (*n* = 6 mice per group). Created with BioRender.com. Visualized using Cytoscape and its CluGo plugin.

At 4 dpi, all stimulation patterns induced a strong inflammatory transcriptional response, together with an upregulation of pathways associated with positive regulation of cell death. Given the extensive neuronal and glial cell death that characterizes the secondary phase of SCI, this response may reflect the persistent injury-related cellular stress at this early stage [44]. All patterns also shared downregulation of neuronal and synaptic processes, including synaptic signaling, action potential generation and ion transport (Fig. 3). Despite this common inflammatory signature, each pattern was associated with a distinct immune-related profile: 10 Hz was associated with myeloid cell differentiation, iTBS with cytokine production and leukocyte activation, and BHFS with leukocyte proliferation and migration. Both iTBS and BHFS were associated with pathways related to phagocytosis, while BHFS showed a stronger association with cytokine response. Notably, BHFS additionally enriched pathways related to tissue repair, including wound healing, fibroblast proliferation and platelet activation (Fig. 3). GSEA further supported these pattern-specific enrichments of inflammatory and tissue-remodeling processes (Figs. S1, S3, S5). Moreover, the GSEA network revealed greater connectivity between adaptive and innate immune response clusters following BHFS, indicating a broader interconnection of immune-related processes at this stage (Fig. S5).

By the end of the treatment at 15 dpi, the transcriptional profiles had diverged substantially between stimulation patterns (Fig. 4). The 10 Hz pattern maintained a predominantly inflammatory and stress-related profile, characterized by persistent inflammatory and apoptotic processes as well as altered reactive oxygen species (ROS) metabolism. Defense responses to virus and interferon-γ-related pathways were downregulated, while GSEA confirmed sustained alterations in inflammatory and synaptic processes (Figs. 4, S2). In contrast, iTBS was associated with a marked upregulation of neuronal-related processes, including synaptic signaling, ion transport and nervous system functions in GO analysis, as well as myelination in GSEA. Downregulated pathways were mainly related in morphogenesis and vascular development, while GSEA also identified a downregulation of stem cell proliferation (Figs. 4, S4). BHFS similarly showed a shift toward neuronal and tissue-repair-related processes, with upregulation of neuronal functions such as ion transport and locomotor rhythm, and downregulation of immune system processes and cytokine-related pathways (Fig. 4). GSEA further identified a downregulation of myelination-related pathways at this stage (Figs. S6).

**Fig. 4.**
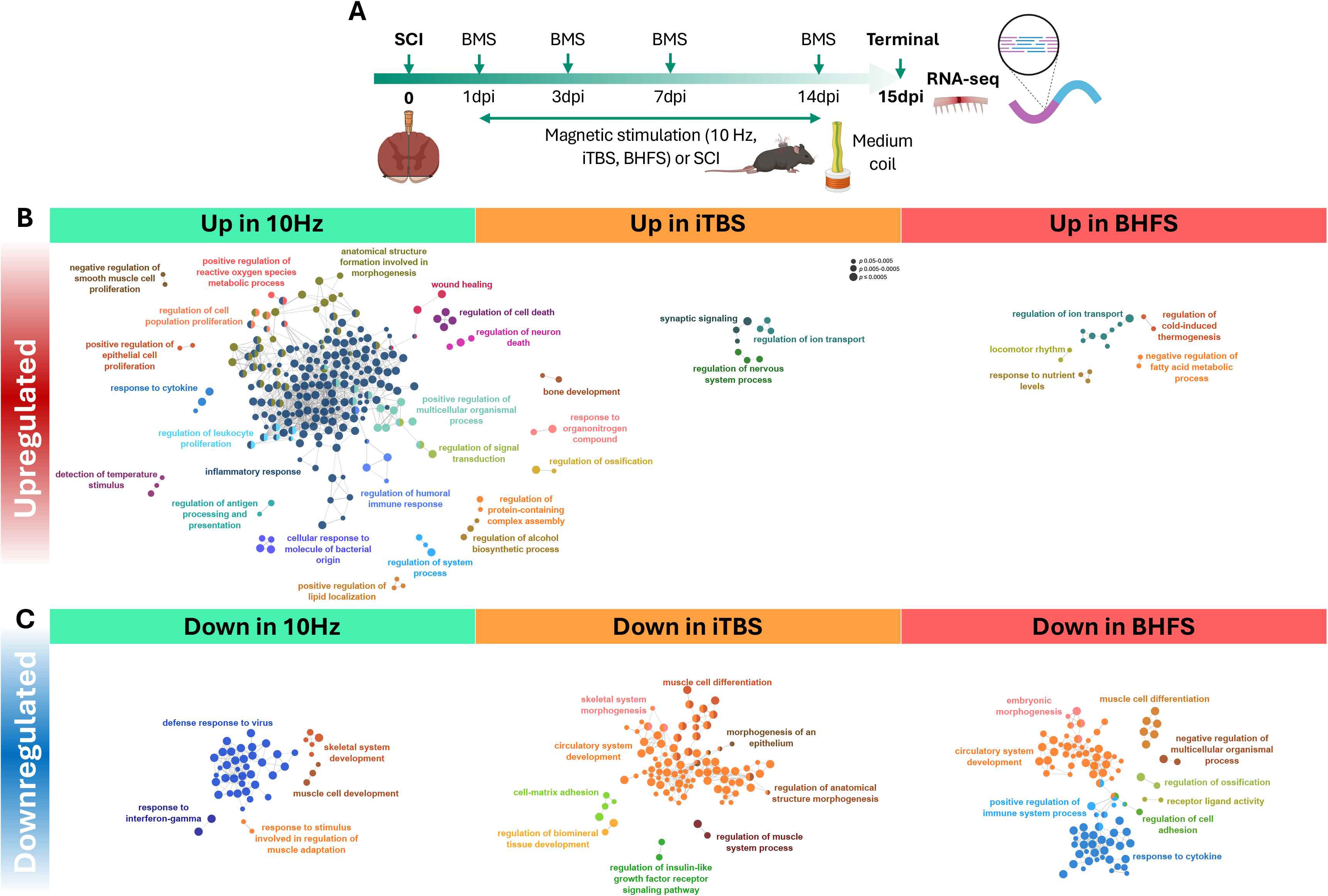
RNA sequencing indicates a persistence of inflammatory signatures under the 10 Hz pattern, whereas iTBS and BHFS promote a return toward normal neuronal expression profiles. **A**. Experimental design: all mice underwent a SCI on day 0, then 3 groups received LI-rTSMS treatment from 1 to 14 dpi using either the 10 Hz, iTBS, or BHFS stimulation pattern, while SCI animals without stimulation served as controls. In parallel, locomotor behaviors were recorded for all mice at 1, 3, 7 and 14 dpi (these behavioral data are presented in Figure 1C), then animals were sacrificed at 15 dpi for RNA-seq analyses. **B–C**. ClueGO was used to identify significantly enriched GO biological processes among upregulated (**B**) and downregulated (**C**) genes in each experimental condition compared to the SCI control. Bubble charts represent enriched GO terms (from left to right: 10 Hz, iTBS, and BHFS), where node size is proportional to the statistical significance of enrichment (*p* values range from ≤ 0.05 to ≤ 0.0005, as indicated by the scale). (*n* = 6 mice per group). Created with BioRender.com. Visualized using Cytoscape and its CluGo plugin.

To further characterize the specific and shared effects of each LI-rTSMS protocol, we performed an overlap analysis of DEGs across the three patterns using Venn diagrams (Figs. S7–S8). After 3 stimulation sessions (4 dpi), 74 genes were downregulated across all three patterns, with a shared GO enrichment restricted to dopamine secretion (Fig. S7). Conversely, 88 genes were upregulated by all three patterns, prominently enriched in GO classes related to inflammation and immune response, such as regulation of phagocytosis, granulocyte migration, and regulation of interleukin-6 production (Fig. S7). Conversely, after 2 weeks treatment (15 dpi), although some overlapping genes persisted, they did not converge on biologically relevant GO classes associated with SCI repair or neural function, further supporting a divergence in the cellular processes modulated by the different stimulation patterns over time (Fig. S8). Pairwise comparisons further revealed that iTBS and BHFS shared a greater proportion of co-regulated genes compared to 10 Hz at both time points (Figs. S7–S8), suggesting that these two complex paradigms engage partially overlapping molecular networks, whereas the 10 Hz pattern triggers a more distinct transcriptional trajectory over time.

Together, these transcriptomic analyses revealed a strong temporal and pattern-dependent modulation of the response to LI-rTSMS, with a common inflammatory signature at 4 dpi followed by increasingly distinct molecular profiles at 15 dpi.

### LI-rTSMS modulates Mi/MΦ distribution and activity at 7 dpi

Based on the locomotor, histological (Figs. 1–2) and RNA-seq findings described above (Figs. 3–4), which revealed marked differences in the inflammatory response and its resolution over time, we therefore investigated whether the differential effects of the three LI-rTSMS patterns were already apparent at an earlier stage following SCI (Fig. 5). Using the same stimulation paradigm, all animals underwent SCI at day 0, and received daily LI-rTSMS treatment or sham stimulation from day 1 to day 6 before being sacrificed for histological analyses the following day (Fig. 5A). The inflammatory response was assessed by evaluating both the total Mi/MΦ population using Iba1, and their activation-associated CD68 expression (Fig. 5B). The positive pixel area of Iba1 and CD68 markers along the rostrocaudal axis was quantified, extending 2.5 mm rostral and caudal to the lesion epicenter. Group comparisons revealed distinct spatial distribution of Iba1 and CD68 following SCI, with CD68 expression being more restricted to the lesion epicenter than Iba1, while both markers displayed pattern-dependent differences in their distribution (Fig. 5B–F). Specifically, the 10 Hz group exhibited a lower Iba1⁺ pixel area from the lesion epicenter toward the caudal region compared with the iTBS and BHFS groups (Fig. 5C), whereas CD68 expression showed an opposite spatial profile at the lesion epicenter, although this difference did not reach statistical significance (Fig. 5D, F).

**Fig. 5.**
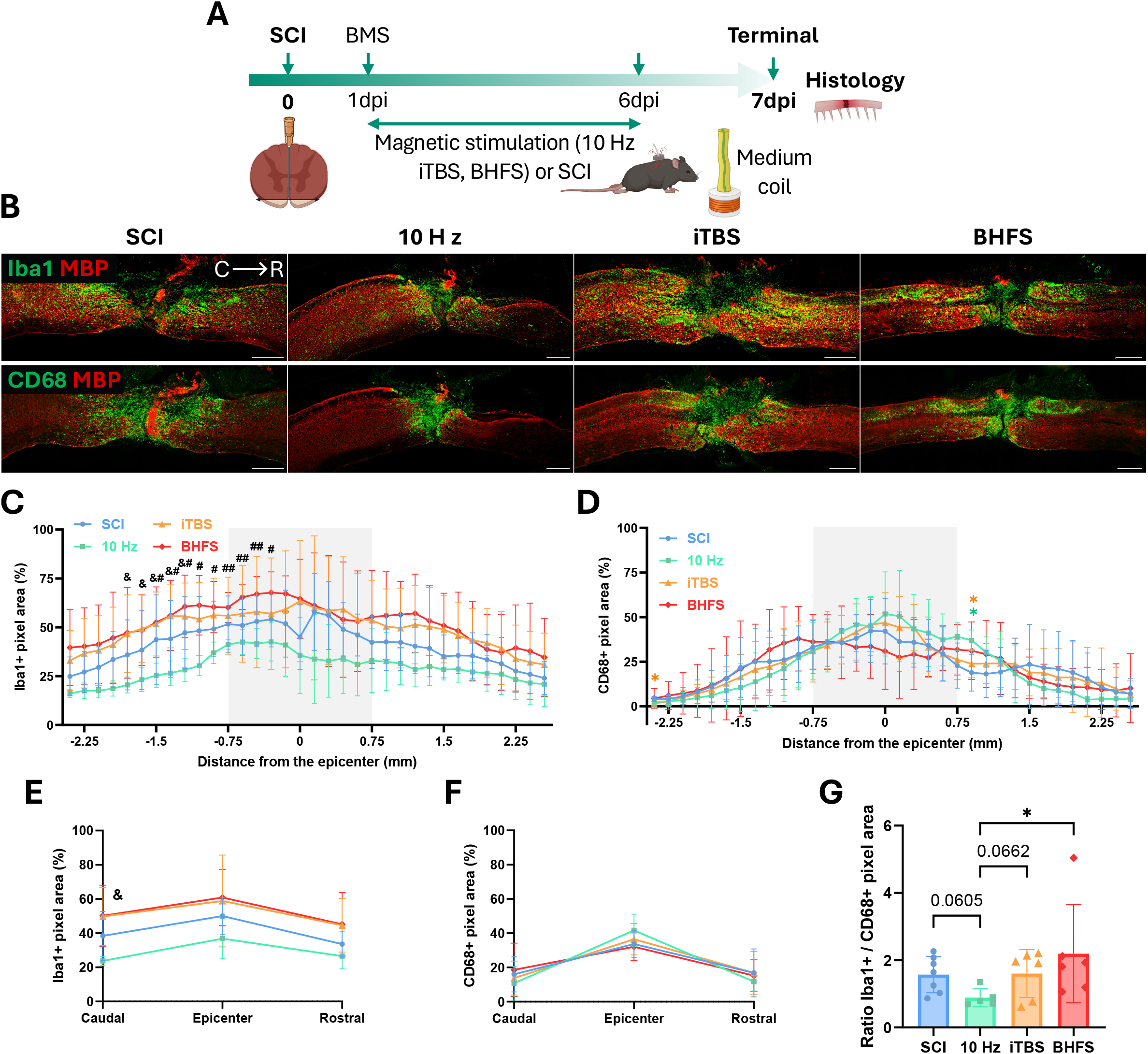
Half the duration of LI-rTSMS treatment modulates Mi/MΦ spatial distribution and activation. **A**. Experimental design: all mice underwent a SCI on day 0, then 3 groups received LI-rTSMS treatment from 1 to 6 dpi, using either the 10 Hz, iTBS, or BHFS stimulation pattern, while SCI animals without stimulation served as controls. The BMS score was assessed at 1 dpi to confirm injury severity consistency across groups. All animals were perfused at 7 dpi for histological analyses. **B–G**. Analysis of the effects of LI-rTSMS on the extent of inflammation and activation state of immune cells between groups. **B**. Representative epifluorescence images (20× magnification) of sagittal spinal cord sections stained with anti-Iba1 (green) and anti-MBP (red) antibodies (top), as well as anti-CD68 (green) and anti-MBP (red) antibodies (bottom), from left to right: SCI, 10 Hz, iTBS, and BHFS. Scale bar: 500 µm. **C**. Quantification of Iba1⁺ pixel area (%) and **D**. CD68⁺ pixel area (%) from -2.4 mm to +2.55 mm around the lesion epicenter. The light grey-shaded areas represent the lesion epicenter. **E**. Quantifications of Iba1⁺ pixel area (%) and **F**. CD68⁺ pixel area (%) across the caudal, epicenter and rostral regions surrounding the lesion. **G**. Ratio measurement of Iba1⁺/CD68⁺ pixel area at the lesion epicenter. Orientation: caudal (C) on the left and rostral (R) on the right in all sagittal sections. Total number of mice: SCI *n* = 7; 10 Hz *n* = 5; iTBS *n* = 6; BHFS *n* = 6. Statistical analyses compared each stimulation group with each other using a Two-Way ANOVA with Tukey’s correction for multiple comparisons (**C– F**), and the Kruskal-Wallis test with trend *p* values indicated on the graph (**G**). (**\*** = *p* ≤ 0.05 between SCI and other groups in their respective color; **&** = *p* ≤ 0.05 between 10 Hz and iTBS; **#** = *p* ≤ 0.05 and **##** = *p* ≤ 0.01 between 10 Hz and BHFS). Created with BioRender.com.

To better illustrate these differences, we calculated an Iba1⁺/CD68⁺ pixel area ratio within the lesion epicenter, defined as the region extending 0.75 mm caudal and rostral to the epicenter. This ratio revealed a distinct shift in the relative distribution of Iba1 and CD68 expression (BHFS: 2.2; 10 Hz: 0.9; SCI and iTBS: 1.6), with a higher value in the BHFS group compared with 10 Hz (Fig. 5G). The lower ratio observed following 10 Hz stimulation therefore indicates relatively greater CD68 expression compared with Iba1 within the lesion.

No significant differences were detected among the groups in Iba1 or CD68 fluorescence intensity, myelin debris area, or MBP colocalization with Iba1 or CD68 (Fig. S9A–H). Furthermore, at this time point, no statistically significant differences were observed between groups in the organization of the fibroglial scar or axon survival/regeneration, although 10 Hz stimulation showed a trend toward increased Tuj1-positive fibers within the lesion (Fig. S9I–L).

These findings indicate that 10 Hz and BHFS patterns induced distinct Mi/MΦ profiles at 7 dpi, with 10 Hz associated with relatively greater CD68 expression compared with Iba1.

#### BHFS enhances proliferation and migration of ependymal cells without changing their differentiation and relative contribution to the glial component of the scar

As previously described, the spinal cord harbors an endogenous stem cell population restricted to ependymal cells surrounding the central canal [17,37]. Several studies have investigated the therapeutic potential of these cells and have therefore sought to modulate the ependymal cell response following SCI [45]. We previously demonstrated that both olfactory ensheathing cell transplantation and high-intensity rTSMS can modulate the proliferation and differentiation of these endogenous stem cells following SCI [27,46].

We therefore investigated the effects of LI-rTSMS on ependymal cell migration, proliferation and differentiation as well as their contribution to glial scar formation (Figs. 6–7). To this end, hFoxJ1- CreERT2::eYFP mice received four daily tamoxifen injections followed by a 10 day washout period before SCI by dorsal hemisection, in order to leave the central canal intact. Half of the animals subsequently received LI-rTSMS treatment for 14 consecutive days, and were sacrificed at 15 dpi for histological analyses. An anti-GFP antibody was used to detect YFP-expressing cells and follow the fate of hFoxJ1-lineage cells (Fig. 6A). Only the BHFS stimulation pattern, which showed the most robust effects in our previous experiments (Figs. 1–5) was used in this study. Transgenic mice that did not undergo SCI were used as non-injured controls (referred as “Control” in Figure 6) to investigate the ependymal cell response compared to SCI groups.

**Fig. 6.**
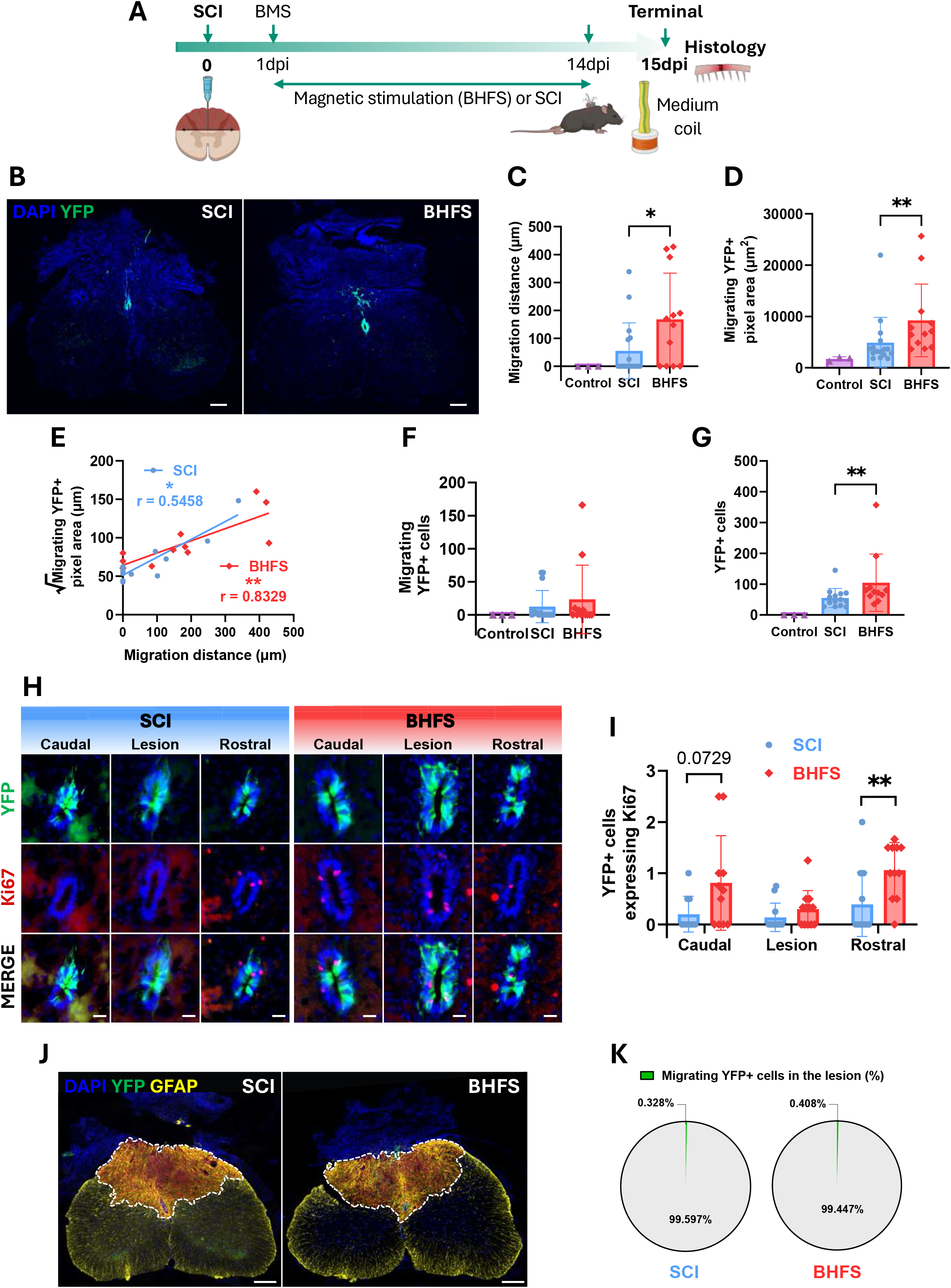
BHFS enhances proliferation and migration of the endogenous ependymal cells toward the injured spinal cord, but does not affect their contribution to the glial scar. **A**. Experimental design: all hFoxJ1-CreERT2::eYFP mice underwent a dorsal hemisection on day 0, half of them received BHFS treatment from 1 to 14 dpi, and animals were perfused at 15 dpi for histological analyses. **B–G**. Images and quantifications of ependymal cell migration into the lesion site. **B**. Representative epifluorescence images (20× magnification) of transverse spinal cord sections stained with DAPI (blue) and anti-GFP (green) antibodies, from left to right: SCI and BHFS. Scale bar: 200 µm. **C**. Analysis of YFP⁺ cells migration distance (µm) (Control *n* = 3; SCI *n* = 17; BHFS *n* = 12) and **D**. their area area occupied within the lesion (µm²) (Control *n* = 3; SCI *n* = 16; BHFS *n* = 12). **E**. Correlation between migration distance (µm) and surface of YFP⁺ cells (µm²) (SCI *n* = 16; BHFS *n* = 12). **F**. Quantification of the number of migrating YFP⁺ cells within the lesion (Control *n* = 3; SCI *n* = 16; BHFS *n* = 12) and **G**. their total number in all the spinal cord (Control *n* = 3; SCI *n* = 14; BHFS *n* = 11). *Note: The uninjured control group is included in panels C, D, F, G for baseline comparison purposes*. **H–I**. Analysis of the proliferation of ependymal cells in the central canal. **H**. Representative epifluorescence images (20× magnification) of transverse spinal cord sections of the central canal from left to right: caudal, lesion epicenter and rostral regions. Sections were stained with DAPI (blue), anti- GFP (green) and anti-Ki67 (red) antibodies, from left to right: SCI and BHFS. Scale bar: 20 µm. **I**. Analysis of YFP⁺ cells expressing Ki67 in each region (SCI *n* = 16; BHFS *n* = 12). **J**. Representative epifluorescence images (20× magnification) of transverse spinal cord sections stained with DAPI (blue), anti-GFP (green) and anti-GFAP (yellow) antibodies. The lesion GFAP-positive is outlined with a white dotted line, and DAPI-positive nuclei within the glial scar are red-labeled, from left to right: SCI and BHFS. Scale bar: 200 µm. **K**. Quantification of the proportion of migrating YFP⁺ cells in the glial scar (%), from left to right: SCI and BHFS. (SCI *n* = 16; BHFS *n* = 12). Statistical analyses were performed using the Mann-Whitney test for pairwise comparisons between groups (**C, D, F, G, I**), and Spearman’s rank correlation and simple linear regression analysis (**E**) (**\*** = *p* ≤ 0.05 and **\*\*** = *p* ≤ 0.01). Created with BioRender.com.

**Fig. 7.**
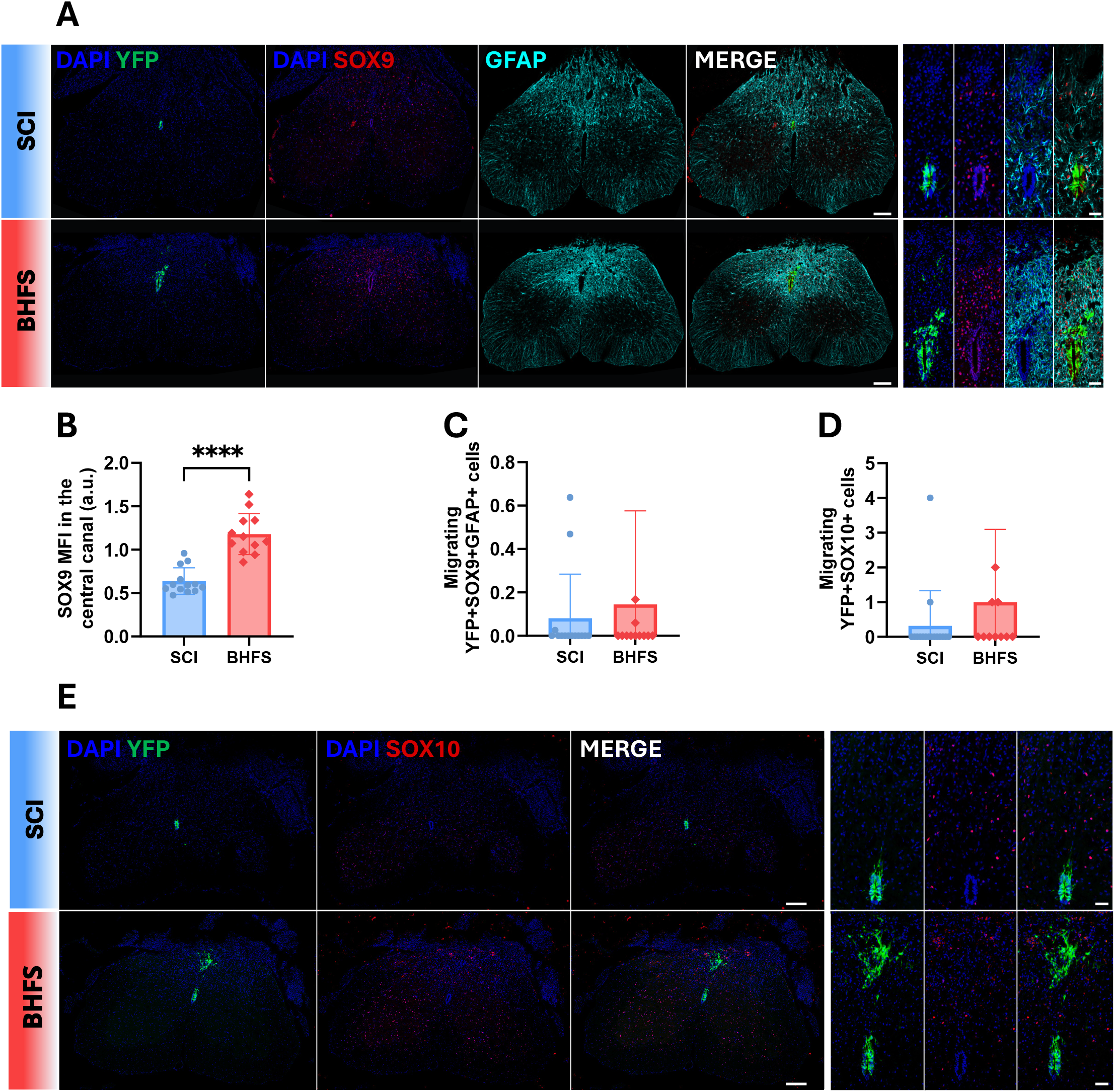
Differentiation of endogenous ependymal cells into astrocytes and oligodendrocytes lineages is not affected by BHFS. **A–E**. Differentiation of ependymal cells into the astrocyte or oligodendrocyte lineage within the dorsal hemisection 15 days post-SCI. **A**. Representative confocal images (20× magnification) of transverse spinal cord sections stained with DAPI (blue), anti-GFAP (cyan), anti-GFP (green) and anti-SOX9 (red) antibodies illustrating astrocyte lineage commitment, from top to bottom: SCI and BHFS. Scale bar: 200 µm. On the right panel: zoom on the central canal. Scale bar: 50 µm. **B**. Quantification of SOX9 MFI in the central canal (SCI *n* = 13; BHFS *n* = 12). **C**. Number of migrating YFP⁺SOX9⁺GFAP⁺ astroglial cells within the lesion (SCI *n* = 14; BHFS *n* = 12). **D**. Number of migrating YFP⁺SOX10⁺ oligodendroglial cells within the lesion (SCI *n* = 16; BHFS *n* = 11). **E**. Representative epifluorescence images (20× magnification) of transverse spinal cord sections stained with DAPI (blue), anti-GFP (green) and anti-SOX10 (red) antibodies, illustrating oligodendrocyte lineage commitment, from top to bottom: SCI and BHFS. Scale bar: 200 µm. On the right panel: zoom on the central canal. Scale bar: 50 µm. Statistical analyses were performed using the Mann-Whitney test for pairwise comparisons between groups (**B–D**) (**\*\*\*\*** = *p* ≤ 0.0001).

We first assessed the impact of the BHFS LI-rTSMS stimulation on the migration of ependymal cells away from the central canal (Fig. 6B–C). We observed that nearly two-thirds of the animals (11 out of 17) in the SCI group exhibited no detectable migration of ependymal-derived cells beyond the central canal (Fig. 6C). In contrast, a greater proportion of mice receiving BHFS LI-rTSMS treatment (8 out of 12) showed cells infiltrating the surrounding parenchyma, with quantitative analysis confirming a significantly increased migration distance compared to SCI alone (Fig. 6C), as well as a larger total area occupied by YFP⁺ cells located outside the central canal (Fig. 6D).

To complement these findings, we assessed the correlation between the maximum migration distance and the area occupied by YFP⁺ cells outside the central canal (Fig. 6E). Interestingly, these two parameters were only weakly correlated in the SCI group (*r = 0.5458*), whereas a strong positive correlation was observed in the BHFS group (*r = 0.8329*) suggesting a more coordinated migratory response following BHFS treatment.

Finally, to distinguish YFP⁺ cells remaining within the central canal from those that had migrated into the spinal parenchyma, we quantified both the number of migrating YFP⁺ cells infiltrating the lesion (Fig. 6F) and the total number of YFP⁺ cells throughout the spinal cord (Fig. 6G). The number of migrating YFP⁺ cells did not significantly differ following BHFS treatment compared to SCI controls (Fig. 6F). Conversely, the total number of YFP⁺ cells within the spinal cord was significantly higher in BHFS-treated animals than in SCI controls (Fig. 6G), suggesting an enhancement of ependymal cell number.

To determine whether these effects were associated with increased ependymal cell proliferation, we quantified the proportion of YFP⁺ cells expressing Ki67 within the central canal across caudal and rostral perilesional regions, as well as at the lesion epicenter (Fig. 6H). Consistent with previous findings showing that proliferating ependymal cells at 15 dpi were predominantly located in the rostral region adjacent to the lesion site and were virtually absent from the lesion epicenter [47], BHFS increased ependymal cell proliferation in the rostral region adjacent to the lesion site, with a trend toward increased proliferation in the caudal region (*p* = 0.0729), whereas no proliferation was detected within the lesion epicenter itself (Fig. 6I).

Finally, to assess the relative contribution of ependymal-derived cells to the glial scar, we calculated the proportion of YFP⁺ cells within the GFAP-positive area at the lesion site (Fig. 6J). These analyses demonstrated that, although LI-rTSMS increased ependymal cell proliferation and migration profile at 15 dpi, the relative contribution of ependymal-derived cells to glial scar formation was not significantly different between the groups. Moreover, ependymal-derived cells represented only a very minor component of the glial scar, accounting for less than 1 % of the total GFAP-positive scar tissue (Fig. 6K).

We next investigated the effects of LI-rTSMS on ependymal cell differentiation at 15 dpi (Fig. 7). First, we quantified their differentiation into glial progenitors using SOX9, a constitutive marker expressed by ependymal cells (Fig. 7A). LI-rTSMS strongly increased SOX9 intensity of ependymal-derived cells in the central canal (Fig. 7B). We next sought to determine whether these SOX9-positive cells exhibited enhanced differentiation toward an astrocytic phenotype within the lesion area. To address this question, we quantified the number of cells located outside the central canal that were triple-positive for YFP, SOX9, and GFAP, which revealed no statistically significant difference between the two experimental conditions (Fig. 7C).

Previous studies have demonstrated that, following SCI, a small proportion of ependymal cells can differentiate into oligodendroglial lineage cells, including oligodendrocyte precursor and mature oligodendrocytes [17,37]. To determine whether this process could be modulated by LI-rTSMS, we quantified the number of YFP⁺ cells expressing the pan-oligodendroglial marker SOX10 (Fig. 7D–E). This analysis demonstrates that differentiation into the oligodendrocyte lineage is a very rare event, being observed in only 2 out of 16 animals in the SCI group, and 4 out of 11 animals in the BHFS group. These findings also indicate that LI-rTSMS does not alter the occurrence of this phenomenon (Fig. 7D– E).

Altogether, our results show that BHFS enhances proliferation and migration, and modulates differentiation of ependymal cells without changing their relative contribution to the glial component of the scar.

### The spatial extent of stimulation is an important determinant of the biological effects of BHFS

As we demonstrated above (Figs. 1–5), the stimulation pattern plays a major role in determining the effects of LI-rTSMS following SCI. To investigate whether other stimulation parameters also contribute to these effects, we examined the impact of coil size, and consequently the spatial extent of the spinal anatomical area being stimulated. To this end, we tested a newly designed small coil (8 mm; being approximately half the diameter of the coil used above) delivering the BHFS pattern at a similar magnetic field intensity (9 mT; designated hereafter as BHFS-S). We first characterized and compared the magnetic fields generated by both the small coil and the standard medium coil using a solenoid-based measurement system (Fig. S10A–B). This analysis confirmed that both coils generated magnetic fields of comparable peak intensity, but exhibited distinct spatial distributions. Specifically, the medium coil (Fig. S10B) produced a more homogenous magnetic field that extended over a larger area than the small coil (Fig. S10A).

Then, we investigated the biological effects of BHFS-S following SCI. As described for the previous experiments (using the same control cohort as in Figures 1 and 2), treatment was administered for 10 min per day over 14 consecutive days. Locomotor performance was assessed at 1, 3, 7 and 14 dpi, and animals were sacrificed at 15 dpi for histological analyses (Fig. 8).

**Fig. 8.**
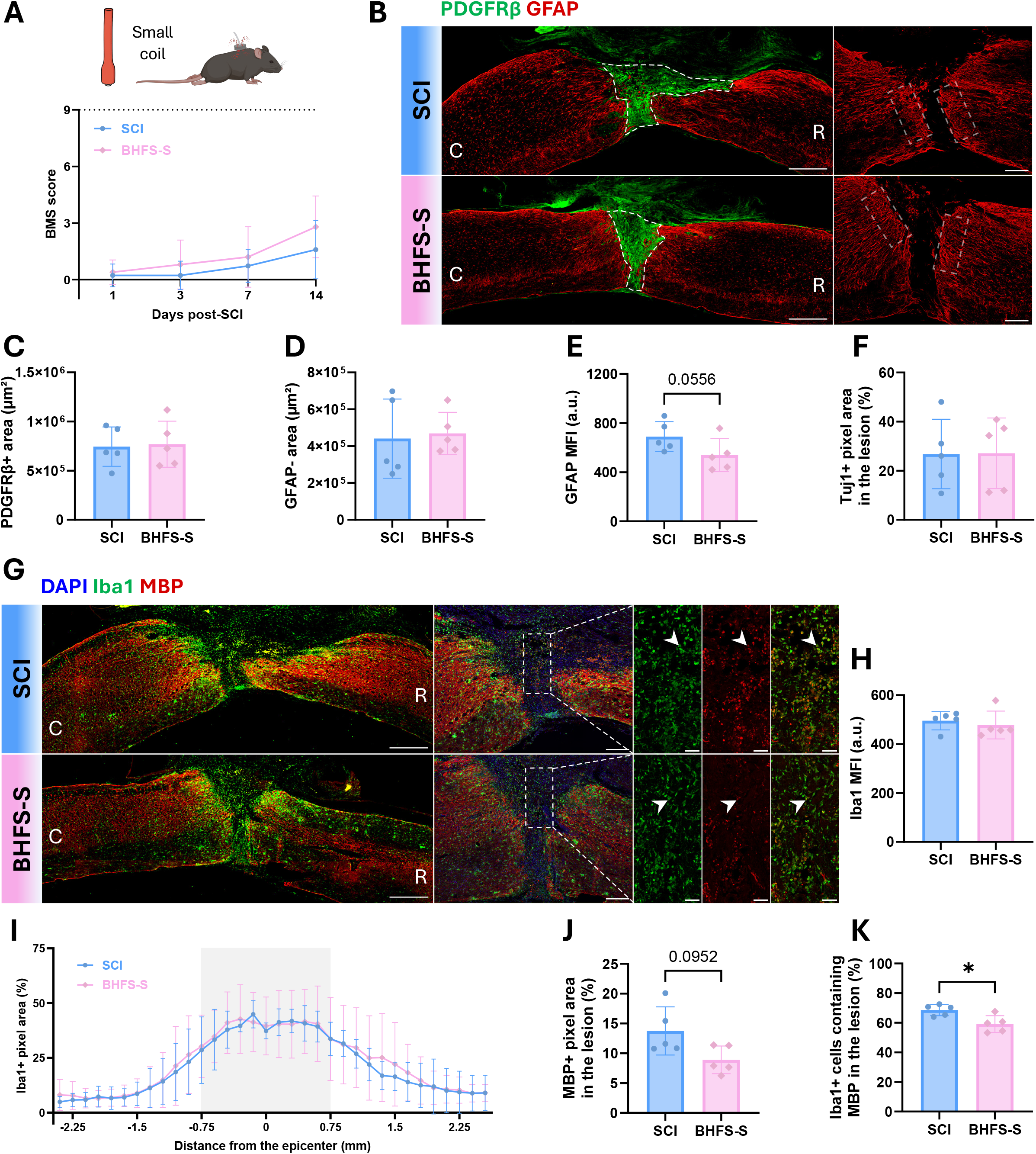
Coil size shapes biological effects using BHFS, and highlights its potential role on myelin debris clearance. **A–K**. All mice underwent a SCI on day 0, then one group received LI-rTSMS treatment, from 1 to 14 dpi, using the small coil and the BHFS pattern (named BHFS-S), while SCI animals without stimulation served as controls. In parallel, locomotor behaviors were recorded for all mice at 1, 3, 7 and 14 dpi, and animals were perfused at 15 dpi for histological analyses. **A**. Assessment of locomotor recovery over time using the BMS score. The white dotted line represents the baseline. **B– E**. Analysis of the effects of LI-rTSMS on fibroglial scar formation. **B**. Representative epifluorescence images (10× magnification) of sagittal spinal cord sections stained with anti-PDGFRβ (green) anti-GFAP (red) antibodies, from top to bottom: SCI and BHFS-S. The white dotted lines surround the lesion core. Scale bar: 500 µm. Right panels show representative confocal images (20× magnification) around the lesion stained with anti-GFAP (red) antibodies, with white dotted lines surrounding the analysed scar borders. Scale bar: 200 µm. **C**. Quantifications of PDGFRβ-positive area (µm²) and **D**. GFAP-negative area (µm²). **E**. Quantification of GFAP MFI at the scar borders. **F**. Analysis of the effects of LI-rTSMS on axonal survival/regrowth within the lesion, represented as Tuj1-positive pixel area within the GFAP- negative lesion area (%). **G–K**. LI-rTSMS effects on inflammation and clearance of myelin debris. **G**. Representative epifluorescence images (10× magnification) of sagittal spinal cord sections stained with DAPI (blue), anti-Iba1 (green) and anti-MBP (red) antibodies, from top to bottom: SCI and BHFS- S. Scale bar: 500 µm. Right panels show representative confocal images (20× magnification). Scale bar: 200 µm. The white dotted rectangle indicates the region zoomed in the panels to the right, with white arrows pointing to colabeled cells. Scale bar: 50 µm. **H**. Quantification of Iba1 MFI. **I**. Quantification of Iba1⁺ pixel area (%) from -2.4 mm to +2.55 mm around the lesion epicenter. The light grey-shaded area represents the lesion epicenter. **J**. Quantification of MBP⁺ pixel area within the MBP-negative lesion area (%) and **K**. the proportion of Iba1⁺ cells containing MBP⁺ debris inside the lesion (%). Orientation: caudal (C) on the left and rostral (R) on the right in all sagittal sections. Statistical analyses were performed using a Mixed-effects analysis with Šidák correction for multiple comparisons (**A**: SCI *n* = 11; BHFS-S *n* = 5), a Mann-Whitney test for pairwise comparisons between groups (**C, D, E, F, H, J, K:** SCI *n* = 5; BHFS-S *n* = 5), and a Two-Way ANOVA with Tukey’s correction for multiple comparisons (**I**). Trend *p* values are indicated on the graphs (**E, J**) (**\*** = *p* ≤ 0.05). Created with BioRender.com.

BHFS-S did not improve locomotor functions, with BMS scores remaining comparable between groups throughout the experimental period (Fig. 8A). Interestingly, whereas BHFS delivered using the medium-sized coil (1.6 cm diameter) previously induced robust modulation of the fibroglial scar (Fig. 1), BHFS-S did not reproduce this effect. Indeed, analysis of PDGFRβ and GFAP immunolabeling revealed no significant differences compared to SCI control in this experimental paradigm (Fig. 8B–D). However, BHFS-S led to a trend toward a decrease in GFAP fluorescence intensity at the scar border (*p = 0.0556*) (Fig. 8B, E). To further characterize the effects of magnetic stimulation on axonal survival/regrowth and Mi/MΦ response, Tuj1 and Iba1 immunolabeling were used, respectively. Neither analysis revealed significant differences between experimental conditions (Fig. 8F–I). BHFS-S showed a trend toward reduced myelin debris area within the lesion (*p = 0.0952*), as assessed by MBP⁺ immunolabeling (Fig. 8J). It also led to a reduction in the proportion of Iba1⁺ cells containing MBP within the lesion (Fig. 8K).

Altogether, these findings demonstrate that coil size, and consequently the spatial extent of the anatomical area exposed to stimulation, is an important determinant of the biological outcomes induced by BHFS following SCI.

### BHFS effects are not associated with detectable neuronal c-Fos induction

While high-intensity rTMS paradigms in the context of depression directly engage specific neuronal populations, such as glutamatergic neurons implied [25,48], whether our LI-rTSMS coils at 9 mT activate spinal neurons remain unknown. We evaluated the immediate-early neuronal activation via c-Fos expression across the entire spinal cord (C1 to L5/L6) following a single session of LI-rTSMS delivered with either the small (BHFS-S) or the medium coil (BHFS-M). Mice were maintained under anesthesia during 1 h after the beginning of the stimulation to eliminate potential confounding behavioral activations such as locomotion or grooming (Fig. 9A).

**Fig. 9.**
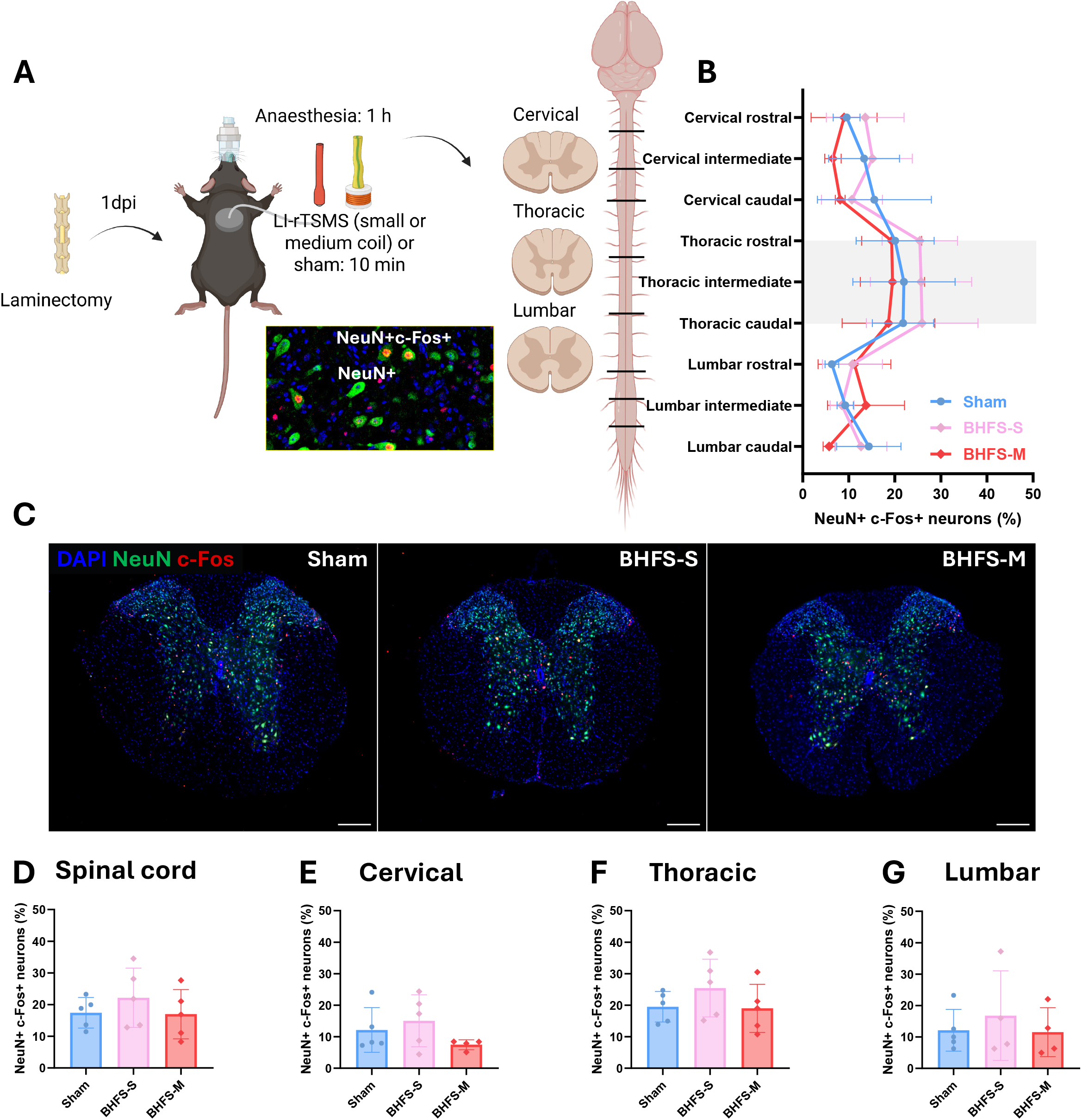
BHFS does not alter neuronal activation in the spinal cord, regardless of coil size. **A–D**. Effect of LI-rTSMS delivered with the small or medium coil on the distribution of neuronal transcriptomic activity in the spinal cord. **A**. Experimental design: all mice underwent a laminectomy on day 0, and were anesthetized using an inhalation mask the next day. Two groups received a single 10 min session of LI-rTSMS treatment using the BHFS pattern delivered with either the small (named BHFS-S) or the medium coil (named BHFS-M), while sham animals without stimulation served as controls. All mice remained under anesthesia for an additional 50 min, resulting in a total anesthesia time of 1 h, before being perfused for histological analyses. Spinal cords were sectioned into cervical, thoracic, and lumbar segments, each subdivided into 3 regions: rostral, intermediate, and caudal. Illustrative epifluorescence image showing co-labelling with DAPI (blue), anti-c-Fos (green) and anti-NeuN (red) antibodies. **B**. Quantification of the proportion of NeuN⁺c-Fos⁺ neurons across all subdivided spinal cord regions. The light grey-shaded area represents the estimated stimulation area. **C**. Representative epifluorescence images (10× magnification) of transverse spinal cord sections stained with DAPI (blue), anti-c-Fos (green) and anti-NeuN (red) antibodies, from left to right: Sham, BHFS-S and BHFS-M. Scale bar: 200 µm. **D–G**. Quantification of the proportion of NeuN⁺c-Fos⁺ neurons in **D**. the entire spinal cord, **E**. the cervical, **F**. the thoracic, and **G**. the lumbar spinal cord segments. Statistical analyses were performed using a Mixed-effect analysis with Dunnett’s correction for multiple comparisons (**B**), and the Kruskal-Wallis test with Dunn’s correction for multiple comparisons (**D–G**). (*n* = 5 mice per group). Created with BioRender.com.

Quantification of the NeuN⁺c-Fos⁺ proportion revealed no significant differences in neuronal activation between non-stimulated controls and mice receiving LI-rTSMS, regardless of the coil size used (Fig. 9B–C). This lack of modulation was consistent across all spatial dimensions: globally throughout the spinal cord (Fig. 9D), when broken down by anatomical region (cervical, thoracic, lumbar; Fig. 9E–G), and when evaluating the dorsal and ventral horns independently (Fig. S10C–J).

These findings demonstrate that the biological effects induced by BHFS on the spinal cord occur independently of detectable neuronal activity in this paradigm.

## Discussion

In our study, we show that LI-rTSMS modulates the injured spinal cord in a pattern- and coil size-dependent manner, with distinct effects on the fibrotic, neural and inflammatory responses to SCI. Notably, the BHFS pattern produced the most pronounced tissue-remodeling effects at 15 dpi, reducing fibrosis, reactive astrogliosis, and myelin debris engulfment by inflammatory cells. These results on phagocytosis were supported by experiments performed with both coil sizes, highlighting the pivotal role of LI-rTSMS on this process. Moreover, distinct inflammatory responses were observed according to the stimulation pattern and the chronology following the lesion. Transcriptomic analyses revealed that LI-rTSMS induced a common early inflammatory response, while each stimulation pattern subsequently produced a distinct molecular signature at later stages. We demonstrated also that LI-rTSMS increased the proliferation and migration of ependymal cells toward the lesion without subsequent detectable differentiation into astrocytes or oligodendrocytes. Finally, to determine whether these effects were associated with early transcriptional changes in neurons, we assessed c-Fos expression in laminectomy-only animals, but LI-rTSMS did not induce any significant changes.

Magnetic stimulation encompasses a wide range of stimulation parameters that differ in frequency, intensity, duration and coil geometry, all of which can influence its biological effects [24,32,49]. Although these parameters have been extensively studied in the brain, their contribution to spinal cord repair remains poorly understood. rTSMS directly targets the injured spinal cord and its surrounding microenvironment, and reported beneficial effects on functional recovery and tissue remodeling [27,50,51]. More recently, Hou *et al.* identified a 40 Hz rTSMS protocol as particularly effective following SCI in promoting locomotor recovery, modifying immune-fibrotic remodeling and enhancing corticospinal tract regeneration [52]. Conversely, another study, using the same frequency parameters reported detrimental behavioral effects following prolonged exposure to a 40 Hz LI-rTMS pattern applied to the brain of young healthy mice, including impaired exploratory behavior and spatial memory [53]. This discrepancy highlights the critical need to better understand frequency- and region-specific effects of repetitive magnetic stimulation, as well as how they vary across different physiological and pathological states of the tissue.

In the present study, we therefore compared three stimulation patterns: 10 Hz, iTBS and BHFS, delivered at a low magnetic field intensity of 9 mT directly over the spinal cord. Our objective was to determine whether low-intensity stimulation patterns could modulate spinal cord repair and whether coil size itself influenced these effects.

First, we performed behavioral assessments using the BMS with the medium-sized coil. At 15 dpi, LI- rTSMS did not significantly improve BMS scores compared with unstimulated SCI animals.

Nevertheless, mean scores remained low (2.38 out of 9), and locomotor recovery had not yet reached a clear plateau at this stage (Fig. 1). Therefore, the absence of a significant functional effect at this time point does not necessarily exclude a later impact of stimulation on neurological recovery.

Then, histological analyses have been performed 15 days post-SCI. We observed pronounced histological changes following BHFS stimulation. The fibroglial scar that forms after injury contributes to the establishment of an extracellular environment that is generally inhibitory to axonal regeneration through the accumulation and altered organization of extracellular matrix components, including chondroitin sulfate proteoglycans, laminin, collagen, and fibronectin [54–56]. BHFS significantly reduced the fibrotic component of the lesion, and decreased GFAP intensity at the scar border, suggesting a beneficial effect on tissue remodeling and a diminution of reactive astrogliosis (Fig. 1) [11,57]. Furthermore, our transcriptomic data at 4 dpi already indicated that BHFS acts on the upregulation of fibroblast proliferation, wound healing and platelet activation (Fig. 3) [58]. Importantly, the reduction of astrogliosis was also observed as a tendency when the smaller coil was used (Fig. 8), indicating that LI-rTSMS may modulate astrocyte responses. Our findings are consistent with previous *in vitro* studies showing that LI-rTMS patterns can modulate astrocyte physiology, hypertrophy, and reactive state, as well as the expression of inflammation- and plasticity-related genes [33,59].

In contrast, LI-rTSMS had limited effects on axonal markers. We observed a trend toward increased Tuj1-positive fibers within the lesion following 10 Hz stimulation at 7 dpi, but this effect was not maintained at a later stage (Figs. 1, S9). This may be related to the lack of resolution of the inflammation observed in this group at 15 dpi. Tuj1 is a neuronal/axonal marker and therefore provides evidence of the presence of neuronal processes within the lesion but does not by itself demonstrate axonal outgrowth or functional reconnection. The absence of a robust effect on Tuj1- positive structures indicates that LI-rTSMS did not produce a detectable increase in neuronal or axonal structures at this time point. However, the absence of improved axonal outgrowth does not exclude the possibility that more prolonged stimulation may result in more pronounced effects on axonal survival and plasticity over time. Overall, these findings suggest that the effects of LI-rTSMS at 15 dpi were more readily detected at the level of the lesion microenvironment than on Tuj1-positive structures [60].

The inflammatory response represents a major component of secondary tissue damage following SCI [61]. Moreover, numerous studies have shown that the beneficial effects of rTMS across various models of neurodegenerative diseases and nervous system injuries are largely mediated by the modulation of the inflammatory response [26,62–64]. Therefore, we investigated the effects of the three stimulation patterns on the inflammatory response. Our results indicate that LI-rTSMS modulates inflammation in a pattern-dependent manner. At 7 dpi, both iTBS and BHFS treatments significantly increased Iba1⁺ pixel area compared to the 10 Hz group across the epicenter to caudal regions, whereas CD68⁺ distribution remained largely comparable among active stimulation groups. However, analysis of the Iba1⁺/CD68⁺ ratio revealed a significant difference between the 10 Hz and BHFS groups, pointing to distinct Mi/MΦ states or densities (Fig. 5). Although Iba1 and CD68 are widely used to characterize inflammatory responses following SCI, these markers are expressed by both resident microglia and infiltrating macrophages. Therefore, our findings should be interpreted as evidence for differences in the overall microglial and macrophage response rather than as selective recruitment of one specific population [58]. Nevertheless, the distinct profiles observed between 10 Hz and BHFS indicate that the stimulation pattern significantly influences the composition of the post-injury inflammatory environment.

This pattern-dependent modulation was further supported by our analysis at 15 dpi. BHFS increases the Mi/MΦ reactivity within the lesion, as assessed by Iba1 fluorescence intensity. It also tends to reduce the amount of myelin debris, and decreases sharply the number of inflammatory cells containing degraded myelin, suggesting lower phagocytic activity at this stage (Fig. 2). Similar effects have recently been reported *in vitro* and *in vivo*, with rTSMS promoting the uptake and clearance of myelin debris by phagocytic cells and implicating the LRP-1 pathway, a receptor involved in this process [31,65]. Furthermore, rTMS has been shown to enhance bactericidal functions in cultured human macrophages through activation of the Nrf2 signaling pathway and inhibition of p38 MAPK, supporting the possibility that magnetic stimulation can modify functional properties of inflammatory cells rather than simply altering their abundance [43]. Importantly, these effects on myelin debris clearance and phagocytosis were also observed using the smaller coil, confirming the involvement of LI-rTSMS in modulating this biological process (Fig. 8).

These observations were also supported by our transcriptomic analyses (Figs. 3–4). At 4 dpi, all stimulation patterns elicited a strong inflammatory signature, indicating that LI-rTSMS reshapes rather than suppresses the early immune response. Importantly, each pattern was associated with a distinct immune-related transcriptional profile, encompassing processes related to myeloid cell differentiation, cytokine production, leukocyte recruitment, and phagocytosis (Fig. 3). These observations are consistent with recent transcriptomic studies showing that rTSMS can primarily affect microglial responses during the early phase following SCI [30,66]. Given that microglia and other glial cells are among the first responders to central nervous system injuries, these findings raise the possibility that LI-rTSMS may be particularly effective when applied early after SCI, as in the present study. Early intervention could allow LI-rTSMS to modulate the initial inflammatory response while maintaining physiological functions such as debris clearance and phagocytosis, potentially supporting tissue repair. Thus, rather than broadly suppressing inflammation, LI-rTSMS may help regulate the early immune response during a critical period of tissue repair. Furthermore, BHFS was the only pattern associated with enrichment of MAPK-related transcriptional programs. MAPK signaling is involved in multiple processes relevant to SCI, including the regulation of inflammatory responses, cell survival, proliferation, and neuronal differentiation. Exposure to electromagnetic fields has been shown to activate p38 MAPK pathway in rat microglia, and modulate the expression of inflammatory mediators, including IL-1β, TNF-α, and IL-10 [67]. rTMS has also been associated with MAPK signaling in the regulation of neural stem cell proliferation and neuronal differentiation [68]. However, Deramaudt *et al*. found an inhibition of p38 MAPK activation following rTMS, as discussed above, suggesting that the effects of magnetic stimulation may depend on stimulation parameters and cellular context [43]. In our study, the selective enrichment of MAPK-related transcriptional programs following BHFS therefore raises the possibility that MAPK signaling contributes to the pattern-specific modulation of the inflammatory and tissue-repair responses induced by LI-rTSMS. However, the functional relevance of this pathway in response to BHFS remains to be determined.

Moreover, the transcriptomic data demonstrated that the effects of LI-rTSMS evolved substantially between the early and later stages following SCI. At 15 dpi, the transcriptional profiles became increasingly pattern-specific. The 10 Hz pattern showed a persistent upregulation of pathways related to the inflammatory response and ROS metabolism, as well as a downregulation of antiviral defense GO terms (Fig. 4). This enrichment may reflect a continuous cellular stress response, consistent with previous evidence that LI-rTMS can modulate ROS-related signaling [69]. Persistent inflammatory signaling at this stage could reflect delayed resolution of the inflammatory response, which is normally expected to progressively transition toward a more reparative environment during the subacute phase following SCI [66]. This observation is particularly relevant given the distinct inflammatory phenotype observed after 10 Hz stimulation at 7 dpi, and may provide a molecular explanation for the absence of a clear tissue-repair benefit associated with this pattern. Furthermore, at 15 dpi, we observed that 40% of the animals in the 10 Hz group (2 out of 5) exhibited extensive spinal cord fibrosis, associated with marked microglial/macrophage infiltration, particularly in the distal region (Figs. 1 and 2). Together, these data suggest that, under the condition tested in the present study, 10Hz stimulation pattern may exacerbate inflammatory and cytotoxicity following SCI, potentially limiting the transition toward tissue repair. Conversely, iTBS and BHFS induced the downregulation of pathways associated with circulatory system development and anatomical structure morphogenesis. This may reflect a shift in the vascular remodeling response, whether these transcriptional changes correspond to reduced vascular remodeling or increased tissue stabilization would require direct assessment of vascular density and maturation [70]. GSEA further identified pattern-specific changes, with iTBS showing downregulation of pathways related to stem cell proliferation, and enrichment of pathways associated with central nervous system myelination, although these transcriptional changes were not accompanied by similarly pronounced tissue-remodeling effects. In contrast, BHFS additionally induced a marked downregulation of immune-related and cytokine-response pathways, reinforcing the histological observation that this pattern may promote the resolution of inflammation during the subacute phase. GSEA also revealed a downregulation of myelination-related pathways following BHFS. This finding is of particular interest in the context of SCI, as oligodendrocyte precursor cells proliferate during the first two weeks after injury and subsequently contribute to the generation of new oligodendrocytes and axon remyelination, with detectable remyelination reported from around 2–3 weeks post-injury [71,72]. These results therefore raise the possibility that BHFS modulates myelination-related processes during the early repair phase following SCI, although whether this transcriptional signature translates into altered oligodendrocyte maturation or remyelination remains unknown.

Together, the transcriptomic findings at 4 and 15 dpi suggest that LI-rTSMS may modulate distinct biological processes over the course of the repair response, including inflammatory pathways at earlier time points, myelination- and tissue-remodeling-related pathways at later stages.

Conventional, high-intensity rTMS are known to influence neuronal excitability through electromagnetic induction, leading to membrane depolarization, action potential generation and intracellular signaling [73,74]. However, whether these mechanisms can be directly extrapolated to LI-rTSMS remains uncertain. Boyer *et al.* demonstrated that 10 Hz LI-rTMS at 10 mT could directly trigger action potentials in cortical neurons, followed by membrane hyperpolarization and reduced intrinsic excitability [75].

In our study, a single session of BHFS applied on laminectomy-only animals did not induce a robust neuronal activation detectable by c-Fos either across the spinal cord, or according to coil size (Figs. 9, S10). However, it does not exclude more subtle changes in neuronal excitability or intracellular signaling, as c-Fos expression is transient and strongly dependent on cell type and timing [76,77]. Consistently, LI-rTMS has been reported to modify intracellular calcium dynamics without necessarily relying on sustained neuronal depolarization [78].

Transcriptomic analyses further support a phase-dependent modulation of neuronal processes. At 4 dpi, all stimulation patterns were associated with downregulation of pathways related to synaptic signaling, action potential generation and ion transport. Given the contribution of neuronal hyperexcitability and excessive glutamatergic signaling contribute to excitotoxic secondary damage after SCI [44], the downregulation of these neuronal pathways may reflect a transient reduction in molecular programs associated with neuronal excitability during the early phase after injury. This interpretation remains speculative because these data are based on transcriptional signatures rather than direct electrophysiological measurements. Conversely, at 15 dpi, iTBS and BHFS were associated with upregulation of neuronal-related pathways, potentially reflecting a shift toward neuronal and synaptic plasticity during the later phase of recovery. These transcriptomic changes may therefore reflect modulation of neuronal state or plasticity, although whether they translate into changes in neuronal activity remains to be determined.

Beyond the effects of repetitive magnetic stimulation on inflammatory modulation and neuronal activity, several studies have also demonstrated that it can modulate the proliferation and differentiation of endogenous neural cells, both in the brain and spinal cord [27,68,79]. We therefore investigated whether LI-rTSMS could modulate the endogenous ependymal response 15 days following SCI (Figs. 6–7). Using hFoxJ1 lineage tracing, we found that BHFS increased the number of ependymal-derived cells and enhanced their proliferation, as indicated by Ki67 expression in the rostral region of the lesion, which is consistent with literature [47]. This proliferation was accompanied by a marked increase in the distance migrated by ependymal-derived cells toward the lesion, although this did not result in an increased number of cells within the lesion itself. It suggests that BHFS may influence the dynamics of endogenous ependymal cells following SCI, particularly by promoting their proliferation and migration, which could reflect changes in cellular motility or in the lesion microenvironment (Fig. 6) [18]. However, additional experiments would be required to determine whether LI-rTSMS directly affects ependymal cell intrinsic properties or instead indirectly modifies the extracellular environment through its effects on inflammatory or glial cells. Moreover, BHFS did not significantly alter the differentiation of ependymal cells toward astrocytic or oligodendrocytic lineages at the investigated time point. The increased SOX9 intensity observed within the central canal may indicate an alteration of the stemness state of ependymal cells (Fig. 7).

Finally, we investigated whether the biological effects of LI-rTSMS could be influenced by coil size, highlighting the importance of the spatial distribution of the magnetic field. Using the same BHFS stimulation pattern and intensity (9 mT) with two coils differing in size (8 and 16 mm, respectively; Fig. S10) allowed us to assess whether the effects of LI-rTSMS were dependent on coil size. Some effects were maintained, such as the modulation of phagocytosis, whereas others, particularly those affecting the fibroglial scar, appeared to be influenced by coil size (Fig. 8). In contrast, a single session of LI-rTSMS did not induce detectable neuronal c-Fos expression regardless of coil size (Figs. 9, S10). Overall, these findings suggest that the size of the anatomical area exposed to the magnetic field, which is determined in part by coil size, may contribute to the biological effects of LI-rTSMS on spinal cord scar remodeling.

### Limitations

Several limitations of this study should be acknowledged. First, the use of a mouse model presents anatomical differences compared to humans, notably the absence of cystic cavity formation following injury. This emphasizes the need to evaluate LI-rTSMS in larger rodent models, such as rats, which better recapitulate post-traumatic cavitation, to confirm the translational potential and reproducibility of these findings.

In our study, we investigated the effects of low-intensity stimulation delivered using three different stimulation patterns. We showed that LI-rTSMS, particularly when delivered using the medium-sized coil and the BHFS pattern, reduces fibrosis and astrocytic reactivity, while also modulating the inflammatory response and the phagocytic activity of Mi/MΦ cells. However, this treatment did not reach the potential to significantly increase axonal outgrowth or improve functional recovery.

Our previous work showed that high-intensity rTSMS (400 mT) could also modulate the fibroglial scar and inflammatory response, while additionally reducing demyelination and promoting locomotor recovery [27]. These findings may suggest that, in this context, high-intensity stimulation could induce more pronounced effects than low-intensity stimulation. However, the results cannot be directly compared, as the stimulation protocols used in these studies were different. In particular, high-intensity stimulation in the previous study was delivered using an intermittent 10 Hz protocol which differs from both the 10 Hz and iTBS patterns used in the present study, notably in terms of stimulation frequency and pattern. Also, due to the size of the coil used to deliver the high-intensity stimulation, the whole spinal cord and brain would have received stimulation, which could account for the more pronounced effects.

Moreover, LI-rTSMS offers several advantages over high-intensity rTSMS. It allows the use of continuous stimulation like 10 Hz, as well as high-frequency stimulation patterns such as iTBS and BHFS, which are more difficult to reproduce at high-intensity because of coil heating and the increased risk of seizures [80–82].

Finally, another limitation of our study is that, although it demonstrates pattern- and coil-dependent responses and points to a central role for the regulation of inflammation in the main effects of LI- rTSMS, it does not precisely define the key mechanisms underlying these effects. Several studies have sought to better understand how rTMS can modulate central nervous system functions, particularly in the context of neurological and psychiatric disorders [25,32,48,62]. Indeed, rTMS has been shown to be dependent on regulatory T cells in the context of Parkinson’s disease [62], other studies have demonstrated that these effects are mediated by a specific population of glutamatergic neurons in the context of depression [25,48]. However, to our knowledge, the mechanisms underlying the effects of rTSMS have not yet been established. Further mechanistic studies are therefore needed to elucidate these mechanisms.

## Conclusion

To conclude, our results demonstrate that LI-rTSMS can modulate the injured spinal cord, particularly by altering the fibroglial scar and inflammatory environment, with no detectable increases in axonal survival/regrowth or functional recovery. These effects on tissues were accompanied by transcriptional changes in pathways related to inflammatory, neuronal, and tissue-remodeling processes, suggesting that LI-rTSMS acts through the coordinated modulation of multiple biological responses after SCI.

Importantly, these effects are strongly influenced by the stimulation pattern used and by coil size, with coil size influencing the spatial distribution and extent of the magnetic field within the spinal cord. This approach opens up new possibilities for future combinatorial therapeutic approaches. In particular, brain and spinal cord stimulation could potentially be combined to simultaneously modulate the lesion environment and promote axonal survival/regrowth of descending pathways. Such a non-invasive therapeutic strategy could bring us one step closer to the clinical translation of rTSMS-based approaches for SCI in humans.

## Acknowledgements

The authors acknowledge the BioMedTech Facilities (INSERM US 36 | CNRS UAR 2009 | Université Paris Cité), in particular the animal facility (Daniel Vintar), as well as the SCM Imaging Facility (Fabrice Licata) for access to the microscopes, and the Zeiss LSM 880 confocal microscope funded through the DIM Cerveau & Pensée and Région Île-de-France. We also thank the SUMMIT (Maisons des modélisations ingénierie et technologies, Sorbonne Université), particularly Alexandre Guerre for designing the magnetic stimulation coils and stimulators used in this project. We are grateful to Prof. Stéphane Vinit and Dr. Pauline Michel-Flutot for performing the magnetic field mapping of the coils presented in Figure S10.

## Contributions

P.N., R.S., F.N. and N.G. conceptualized the project. P.N., R.S., F.N. and N.G. designed the experiments. P.N., A.D., F.S., L.M., A.F., C.S., A.MG., E.D.M., and C.R. performed the experiments. P.N., A.D., F.S., L.M., A.F., C.S. and A.MG. analyzed the results. P.N., A.D., F.S., R.S., F.N. and N.G. wrote the article.

## Declarations

### Ethics approval and consent to participate

All experimental procedures complied with the European Community guidelines for the care and use of laboratory animals (86/609/EEC; Official Journal of the European Communities No. L358, Decembre 18, 1986), The French Decree No. 87/848 of October 19, 1987. All procedures that involved animal experiments were approved by the local Ethics Committee on January 10^th^ 2024, for a five years period, with approval number of APAFIS #45711-202307121857640 v9. Title of the approved project was “ Synergistic Effects of a Combined Therapy for Spinal Cord Injury Repair: Biomaterial Implants, Recruitment of Endogenous Stem Cells, and Magnetic Stimulation”.

### Competing interests

The authors declare no competing interests.

All authors agree with the submission of this manuscript.

**Table. S1. Parameters used for GO enrichment and network clustering analysis.** Summary of the bioinformatic settings applied to the functional enrichment analysis of DEGs presented in Figures 3–4 and to the Venn diagram analyses in Figures S7–S8. The table details the prefiltering thresholds, ontology database versions, network specificity parameters, advanced term selection criteria, clustering criteria, statistical correction method (Bonferroni step-down), and grouping options, including term fusion, initial group sizes, and merging percentages.

**Table S2: GO terms used in GO analyses, related to Figures 3 and 4**. The file contains 12 sheets: the first lists the GO terms downregulated in 10 Hz group at 4 dpi, the second lists the GO terms upregulated in 10 Hz group at 4 dpi, the third lists the GO terms downregulated in iTBS group at 4 dpi, the fourth lists the GO terms upregulated in iTBS group at 4 dpi, the fifth lists the GO terms downregulated in BHFS group at 4 dpi, the sixth lists the GO terms upregulated in BHFS group at 4 dpi, the seventh lists the GO terms downregulated in 10 Hz group at 15 dpi, the eighth lists the GO terms upregulated in 10 Hz group at 15 dpi, the ninth lists the GO terms downregulated in iTBS group at 15 dpi, the tenth lists the GO terms upregulated in iTBS group at 15 dpi, the eleventh lists the GO terms downregulated in BHFS group at 15 dpi and the twelth lists the GO terms upregulated in BHFS group at 15 dpi.

**Table S3: DEGs and GO terms used in GSEA analysis in 10 Hz vs SCI at 4 and 15 dpi, related to Figures S1 and S2.** The file contains 10 sheets: the first lists the DEGs downregulated at 4 dpi, the second lists the DEGs upregulated at 4 dpi, the third presents the raw analysis of downregulated GO terms at 4 dpi, the fourth presents the raw analysis of upregulated GO terms at 4 dpi, the fifth contains the GO terms that are illustrated in the GSEA representation generated using Cytoscape at 4 dpi, the sixth lists the DEGs downregulated at 15 dpi, the seventh lists the DEGs upregulated at 15 dpi, the eighth presents the raw analysis of downregulated GO terms at 15 dpi, the ninth presents the raw analysis of upregulated GO terms at 15 dpi, the tenth contains the GO terms that are illustrated in the GSEA representation generated using Cytoscape at 15 dpi.

**Table S4: DEGs and GO terms used in GSEA analysis in iTBS vs SCI at 4 and 15 dpi, related to Figures S3 and S4.** The file contains 10 sheets: the first lists the DEGs downregulated at 4 dpi, the second lists the DEGs upregulated at 4 dpi, the third presents the raw analysis of downregulated GO terms at 4 dpi, the fourth presents the raw analysis of upregulated GO terms at 4 dpi, the fifth contains the GO terms that are illustrated in the GSEA representation generated using Cytoscape at 4 dpi, the sixth lists the DEGs downregulated at 15 dpi, the seventh lists the DEGs upregulated at 15 dpi, the eighth presents the raw analysis of downregulated GO terms at 15 dpi, the ninth presents the raw analysis of upregulated GO terms at 15 dpi, the tenth contains the GO terms that are illustrated in the GSEA representation generated using Cytoscape at 15 dpi.

**Table S5: DEGs and GO terms used in GSEA analysis in BHFS vs SCI at 4 and 15 dpi, related to Figures S5 and S6.** The file contains 10 sheets: the first lists the DEGs downregulated at 4 dpi, the second lists the DEGs upregulated at 4 dpi, the third presents the raw analysis of downregulated GO terms at 4 dpi, the fourth presents the raw analysis of upregulated GO terms at 4 dpi, the fifth contains the GO terms that are illustrated in the GSEA representation generated using Cytoscape at 4 dpi, the sixth lists the DEGs downregulated at 15 dpi, the seventh lists the DEGs upregulated at 15 dpi, the eighth presents the raw analysis of downregulated GO terms at 15 dpi, the ninth presents the raw analysis of upregulated GO terms at 15 dpi, the tenth contains the GO terms that are illustrated in the GSEA representation generated using Cytoscape at 15 dpi.

**Fig. S1. GSEA and differentially enriched biological processes in the 10 Hz-treated group at 4 dpi. A–B**. GSEA is represented as bubble charts of regulated GO terms ("Biological Process" category) in the 10 Hz-treated group compared to the SCI control after 3 days of LI-rTSMS, and perfusion for RNA-seq analysis at 4 dpi. These GO terms are **A**. downregulated (blue panel) or **B**. upregulated (pink panel). Each dot represents a regulated GO term; their size and color intensity are proportional to the number of genes associated with the GO term (ranging from 0 to 200) and the enrichment adjusted *p* value (false discovery rate *q* value), respectively. GO terms were grouped into categories using the AutoAnnotate plugin (*n* = 6 mice per group). Visualized using Cytoscape.

**Fig. S2. GSEA and differentially enriched biological processes in the 10 Hz-treated group at 15 dpi. A–B**. GSEA is represented as bubble charts of regulated GO terms ("Biological Process" category) in the 10 Hz-treated group compared to the SCI control after 14 days of LI-rTSMS, and perfusion for RNA-seq analysis at 15 dpi. These GO terms are **A**. downregulated (blue panel) or **B**. upregulated (pink panel). Each dot represents a regulated GO term; their size and color intensity are proportional to the number of genes associated with the GO term (ranging from 0 to 200) and the enrichment adjusted *p* value (false discovery rate *q* value), respectively. GO terms were grouped into categories using the AutoAnnotate plugin (*n* = 6 mice per group). Visualized using Cytoscape.

**Fig. S3. GSEA and differentially enriched biological processes in the iTBS-treated group at 4 dpi. A–B**. GSEA is represented as bubble charts of regulated GO terms ("Biological Process" category) in the iTBS-treated group compared to the SCI control after 3 days of LI-rTSMS, and perfusion for RNA-seq analysis at 4 dpi. These GO terms are **A**. downregulated (blue panel) or **B**. upregulated (pink panel). Each dot represents a regulated GO term; their size and color intensity are proportional to the number of genes associated with the GO term (ranging from 0 to 200) and the enrichment adjusted *p* value (false discovery rate *q* value), respectively. GO terms were grouped into categories using the AutoAnnotate plugin (*n* = 6 mice per group). Visualized using Cytoscape.

**Fig. S4. GSEA and differentially enriched biological processes in the iTBS-treated group at 15 dpi. A–B**. GSEA is represented as bubble chartsof regulated GO terms ("Biological Process" category) in the iTBS-treated group compared to the SCI control after 14 days of LI-rTSMS, and perfusion for RNA-seq analysis at 15 dpi. These GO terms are **A**. downregulated (blue panel) or **B**. upregulated (pink panel). Each dot represents a regulated GO term; their size and color intensity are proportional to the number of genes associated with the GO term (ranging from 0 to 200) and the enrichment adjusted *p* value (false discovery rate *q* value), respectively. GO terms were grouped into categories using the AutoAnnotate plugin (*n* = 6 mice per group). Visualized using Cytoscape.

**Fig. S5. GSEA and differentially enriched biological processes in the BHFS-treated group at 4 dpi. A–B**. GSEA is represented as bubble charts of regulated GO terms ("Biological Process" category) in the BHFS-treated group compared to the SCI control after 3 days of LI-rTSMS, and perfusion for RNA-seq analysis at 4 dpi. These GO terms are **A**. downregulated (blue panel) or **B**. upregulated (pink panel). Each dot represents a regulated GO term; their size and color intensity are proportional to the number of genes associated with the GO term (ranging from 0 to 200) and the enrichment adjusted *p* value (false discovery rate *q* value), respectively. GO terms were grouped into categories using the AutoAnnotate plugin (*n* = 6 mice per group). Visualized using Cytoscape.

**Fig. S6. GSEA and differentially enriched biological processes in the BHFS-treated group at 15 dpi. A–B**. GSEA is represented as bubble charts of regulated GO terms ("Biological Process" category) in the BHFS-treated group compared to the SCI control after 14 days of LI-rTSMS, and perfusion for RNA-seq analysis at 15 dpi. These GO terms are **A**. downregulated (blue panel) or **B**. upregulated (pink panel). Each dot represents a regulated GO term; their size and color intensity are proportional to the number of genes associated with the GO term (ranging from 0 to 200) and the enrichment adjusted *p* value (false discovery rate *q* value), respectively. GO terms were grouped into categories using the AutoAnnotate plugin (*n* = 6 mice per group). Visualized using Cytoscape.

**Fig. S7. Venn diagrams of GO terms comparisons between LI-rTSMS patterns at 4 dpi. A–B**. LI-rTSMS groups compared to the SCI control after 3 days of treatment, and perfusion for RNA-seq analysis at 4 dpi. Venn diagrams show the overlap of the GO terms which are **A**. upregulated (top) or **B**. downregulated (bottom) among the three LI-rTSMS treatment groups: 10 Hz (green), iTBS (red), and BHFS (blue). The non-overlapping regions represent biological processes specifically regulated by each stimulation pattern, while the overlapping regions identify clusters commonly shared between two or all three stimulation protocols. (*n* = 6 mice per group). Visualized using Cytoscape.

**Fig. S8. Venn diagrams of GO terms comparisons between LI-rTSMS patterns at 15 dpi. A–B**. LI-rTSMS groups compared to the SCI control after 14 days of treatment, and perfusion for RNA-seq analysis at 15 dpi. Venn diagrams show the overlap of the GO terms which are **A**. upregulated (top) or **B**. downregulated (bottom) among the three LI-rTSMS treatment groups: 10 Hz (green), iTBS (red), and BHFS (blue). The non-overlapping regions represent biological processes specifically regulated by each stimulation pattern, while the overlapping regions identify clusters commonly shared between two or all three stimulation protocols. (*n* = 6 mice per group). Visualized using Cytoscape.

**Fig. S9. Evaluation of half-duration LI-rTSMS effects on inflammation, fibroglial scar, and axonal features. A–H**. LI-rTSMS effects on inflammation and clearance of myelin debris at 7 days post-SCI. **A**. Illustrative confocal image (20× magnification) of a sagittal spinal cord section stained with DAPI (blue), anti-Iba1 (green) and anti-MBP (red) antibodies from an iTBS-treated mouse. Scale bar: 200 µm. The white dotted rectangle indicates the region zoomed in the panels to the right, with white arrows pointing to colabeled cells. Scale bar: 50 µm. **B**. Illustrative confocal image (20× magnification) of a sagittal spinal cord section stained with DAPI (blue), anti-CD68 (green) and anti-MBP (red) antibodies from an iTBS-treated mouse. Scale bar: 200 µm. The white dotted rectangle indicates the region zoomed in the panels to the right, with white arrows pointing to colabeled cells. Scale bar: 50 µm. **C**. Quantification of Iba1 MFI, and **D**. CD68 MFI. **E**. Quantification of MBP-negative area (µm²), and **F**. the MBP⁺ pixel area within this lesion area (%). **G**. Quantification of the proportion of Iba1⁺ cells containing MBP⁺ debris inside the lesion (%), and **H**. the proportion of CD68⁺ cells containing MBP⁺ debris inside the lesion (%). **I–L**. Analysis of the effects of LI-rTSMS on fibroglial scar formation and axonal survival/regrowth. **I**. Representative epifluorescence images (10× magnification) of sagittal spinal cord sections stained with anti-PDGFRβ (green) and anti-GFAP (red) antibodies (top) as well as anti-Tuj1 (cyan) and anti-GFAP (red) antibodies (bottom), from left to right: SCI, 10 Hz, iTBS, and BHFS. Scale bar: 500 µm. **J**. Quantifications of PDGFRβ-positive area (µm²), **K**. GFAP-negative area (µm²), and **L**. Tuj1-positive pixel area within the GFAP-negative lesion area (%). Orientation: caudal (C) on the left and rostral (R) on the right in all sagittal sections. Total number of mice: SCI *n* = 7; 10 Hz *n* = 5; iTBS *n* = 6; BHFS *n* = 6. Statistical analyses compared the SCI group with each stimulation group using the Kruskal-Wallis test with Dunn’s correction for multiple comparisons. Trend *p* value is indicated on the graph (**L**).

**Fig. S10. Solenoid of small and medium LI-rTSMS coils, and c-Fos mapping of the dorsal and the ventral horns of the spinal cord. A–B**. Three-dimensional distribution of the magnetic field intensity measured using a solenoid and expressed in volt along the X, Y, and Z axes for **A**. the small and **B**. the medium coil. **C–J**. Analyses of the effects of a single session of LI-rTSMS on neuronal activation across the dorsal and ventral horns of the spinal cord. **C–F**. Quantification of the proportion of NeuN⁺c-Fos⁺ neurons in the dorsal horn of **C**. the entire spinal cord, **D**. the cervical, **E**. the thoracic, and **F**. the lumbar spinal cord segments. **G– I**. Quantification of the proportion of NeuN⁺c-Fos⁺ neurons in the ventral horn of **G**. the entire spinal cord, **H**. the cervical, **I**. the thoracic, and **J**. the lumbar spinal cord segments. Statistical analyses compared the SCI group with each stimulation group using the Kruskal-Wallis test with Dunn’s correction for multiple comparisons. (*n* = 5 mice per group). Created with BioRender.com.

