## Supplementary material for "Pattern-dependent low-intensity repetitive magnetic stimulation enhances spinal cord repair through modulation of neuroinflammation": Table S1

| | ALL | VENN DIAGRAM |
|---|---|---|
| **Prefiltering on DEGs** | pAdj ≤ 0.05<br>400 top up or down genes | |
| **Ontologies/Pathways**<br>**Network Specificity**<br>**Use GO Term Fusion**<br>**Show only Pathways with pV ≤ 0.05** | Go-Biological Process 25-05-22<br>Medium<br>Yes<br>Yes | Go-Biological Process 25-05-22<br>Medium<br>Yes<br>Yes |
| **Advanced Term Selection Options** | | |
| **Go Tree interval**<br>**Cluster** | 4<br>3 Min #Genes and 4% Genes | 4<br>3 Min #Genes and 4% Genes |
| **Go Term Network Connectivity: Kappa Score**<br>**Statistical Options** | 0.4<br>Default: Bonferroni step down | 0.4<br>Default: Bonferroni step down |
| **Grouping Options** | | |
| **Use Go Term Grouping**<br>**Leading Group Term based on**<br>**Initial Group Size**<br>**%Genes for Group Merge**<br>**%Term for Group Merge** | Yes<br>Highest Significance<br>2<br>30<br>30 | Yes<br>Highest Significance<br>2<br>50<br>50 |
