## Supplementary figures and images for "Pattern-dependent low-intensity repetitive magnetic stimulation enhances spinal cord repair through modulation of neuroinflammation"

### Supplementary S1

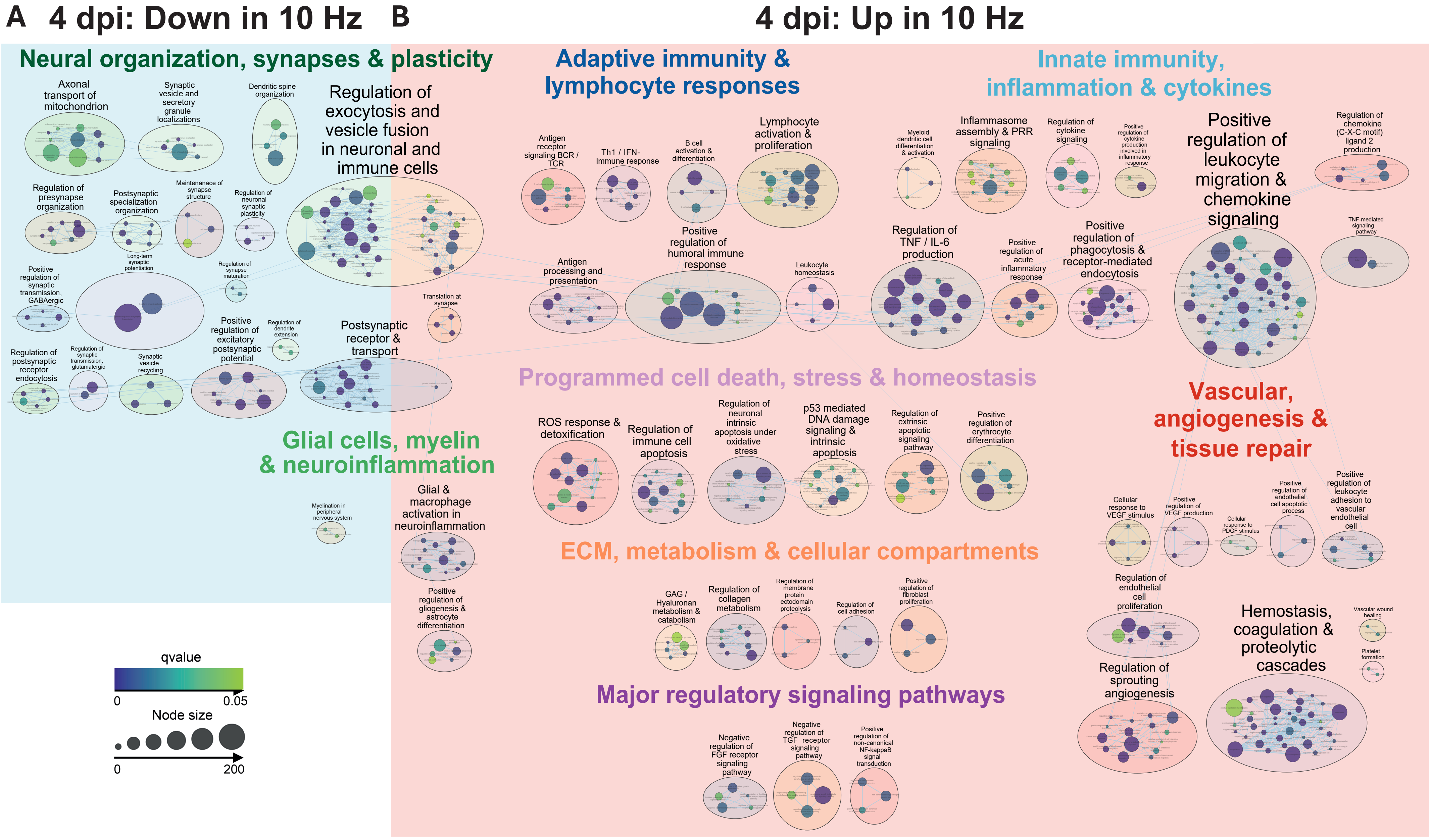

### Supplementary S2

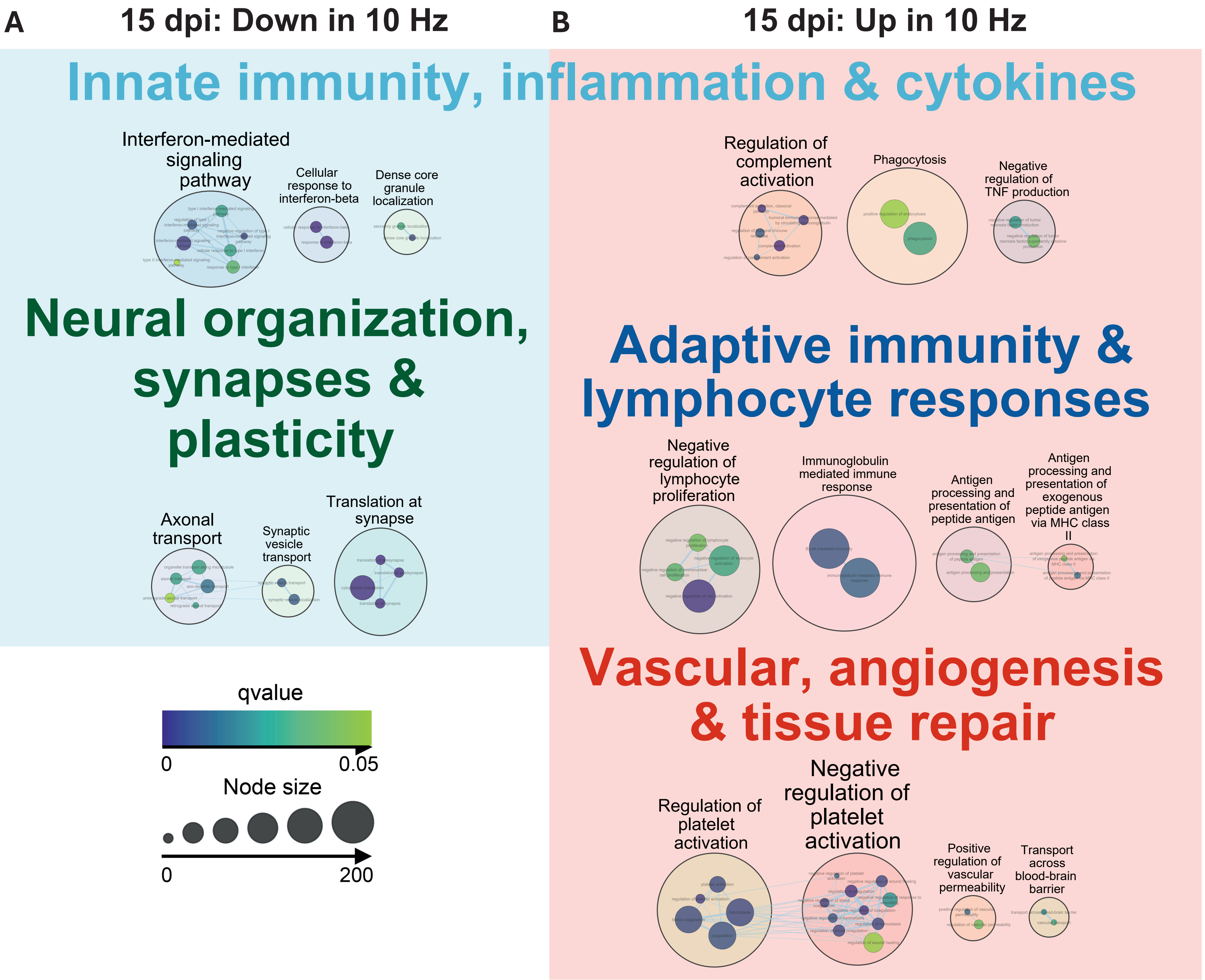

### Supplementary S3

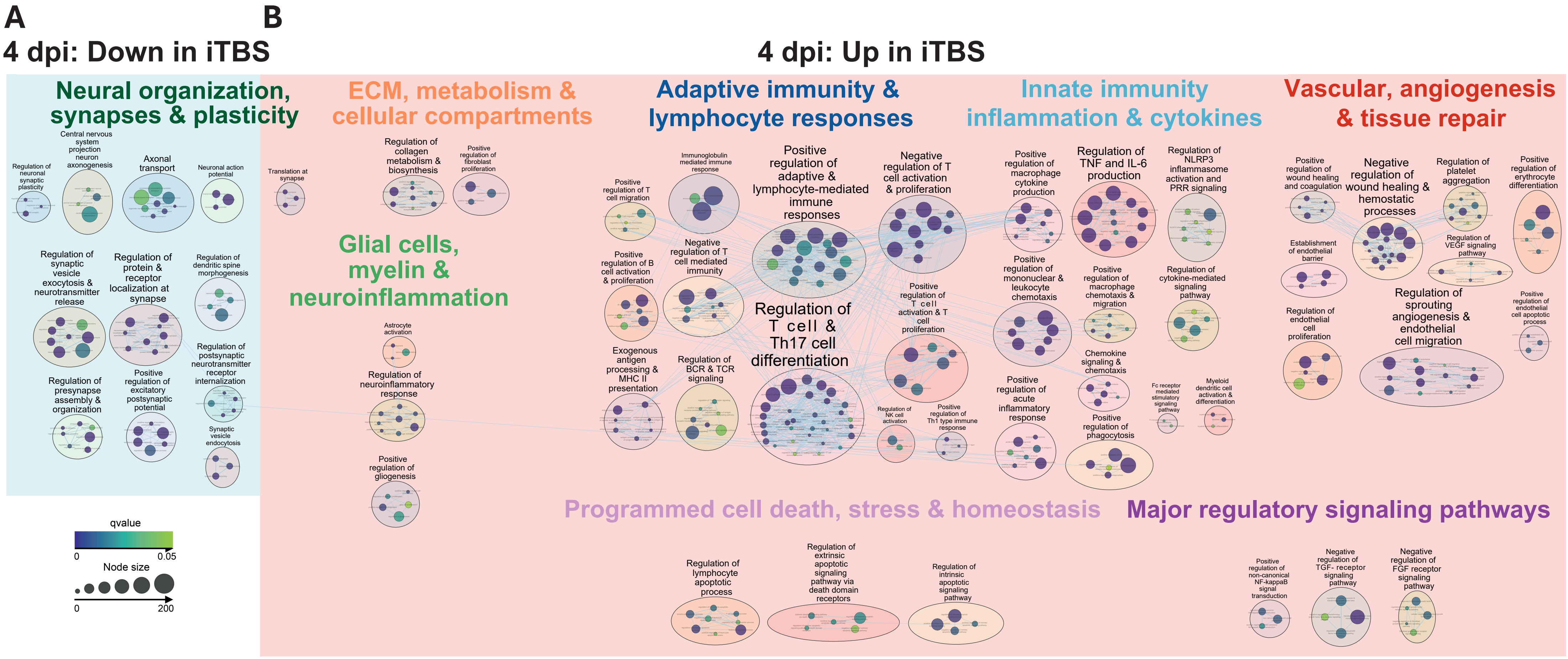

### Supplementary S4

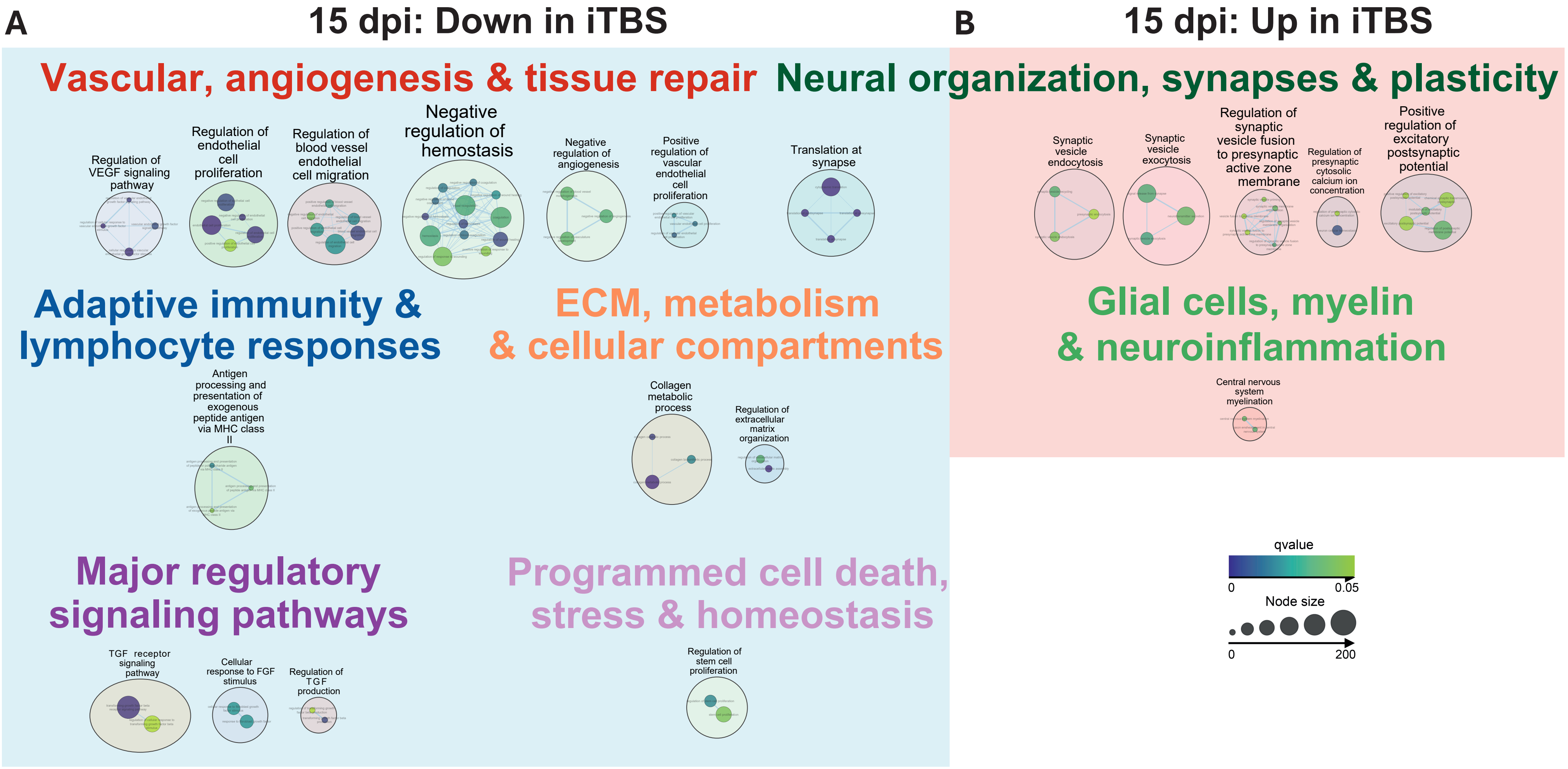

### Supplementary S5

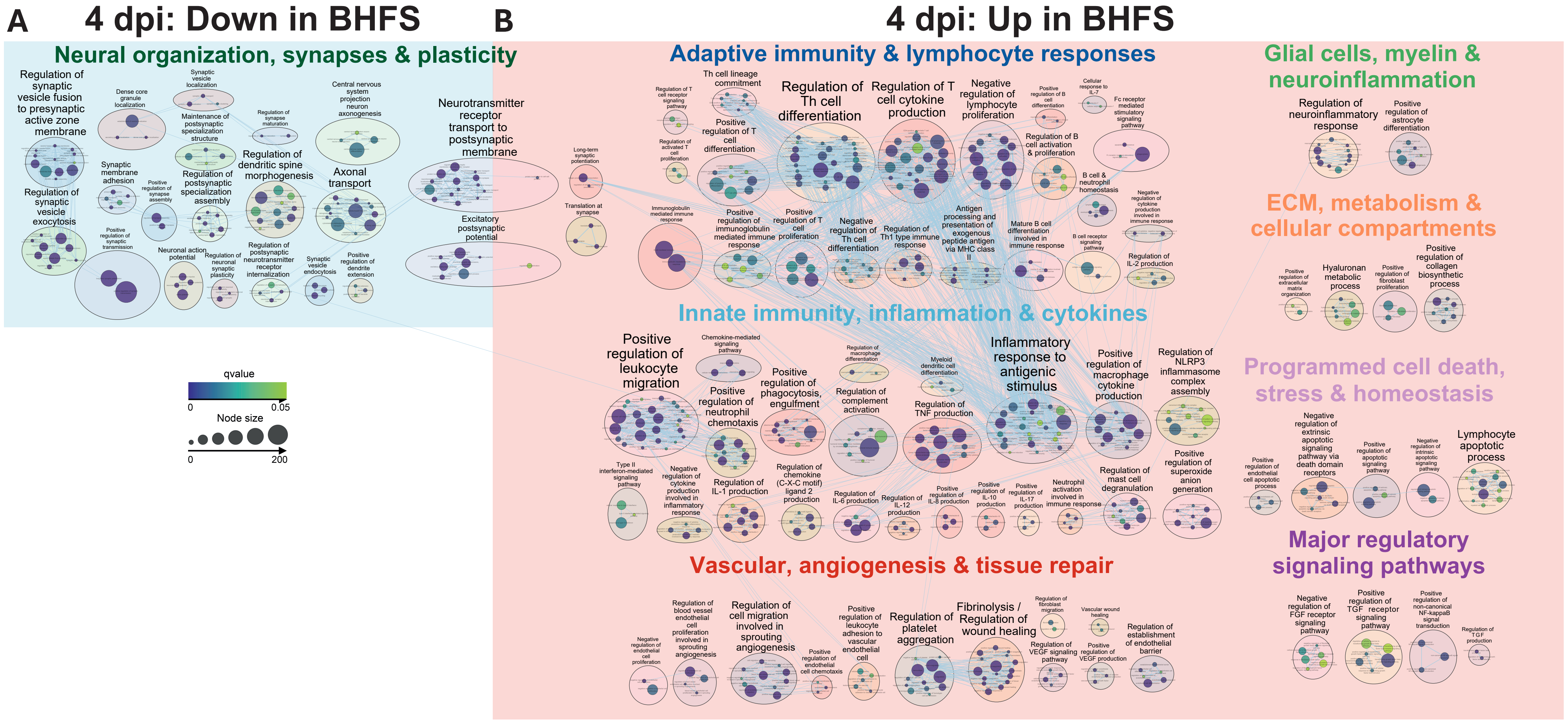

### Supplementary S6

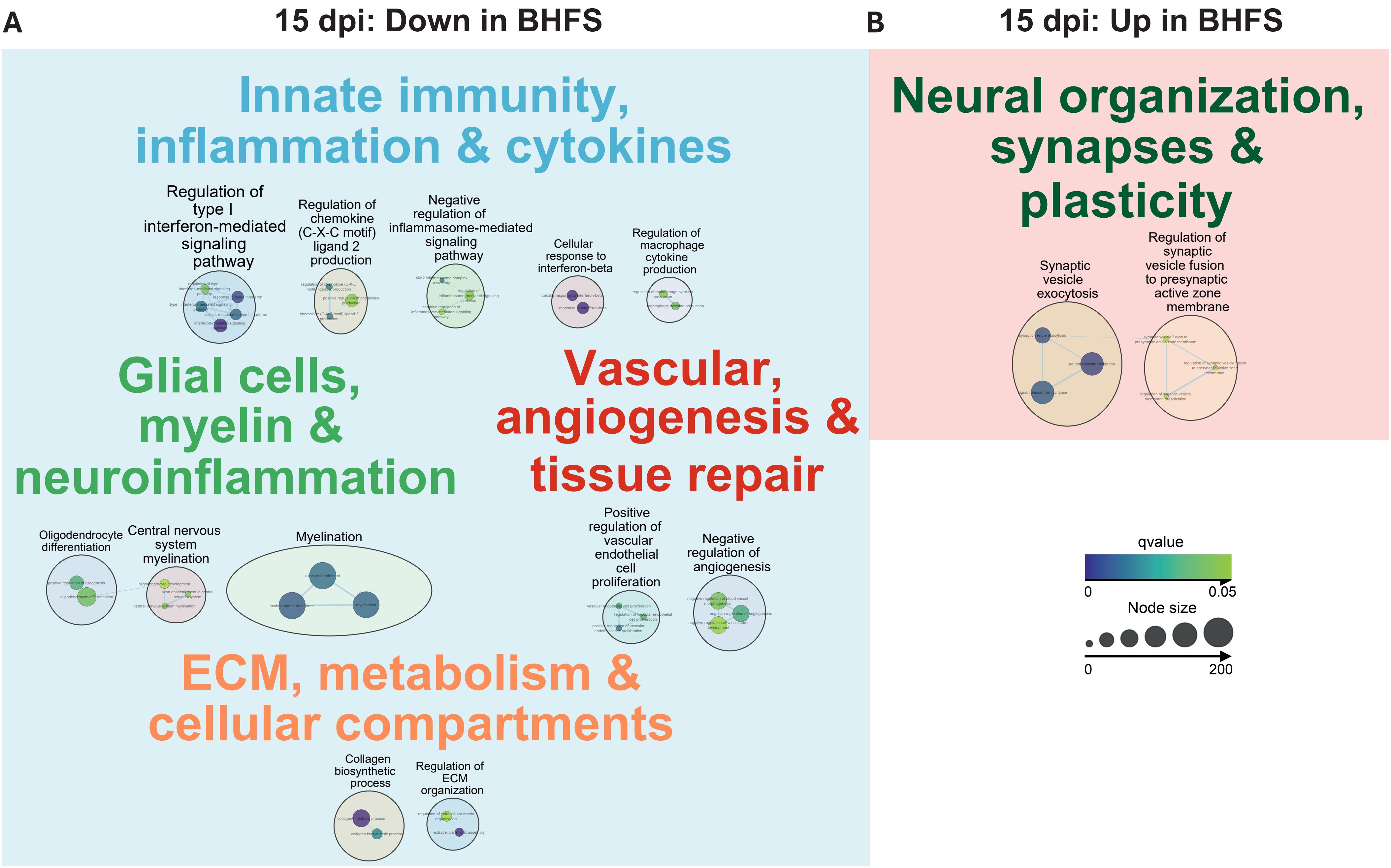

### Supplementary S7

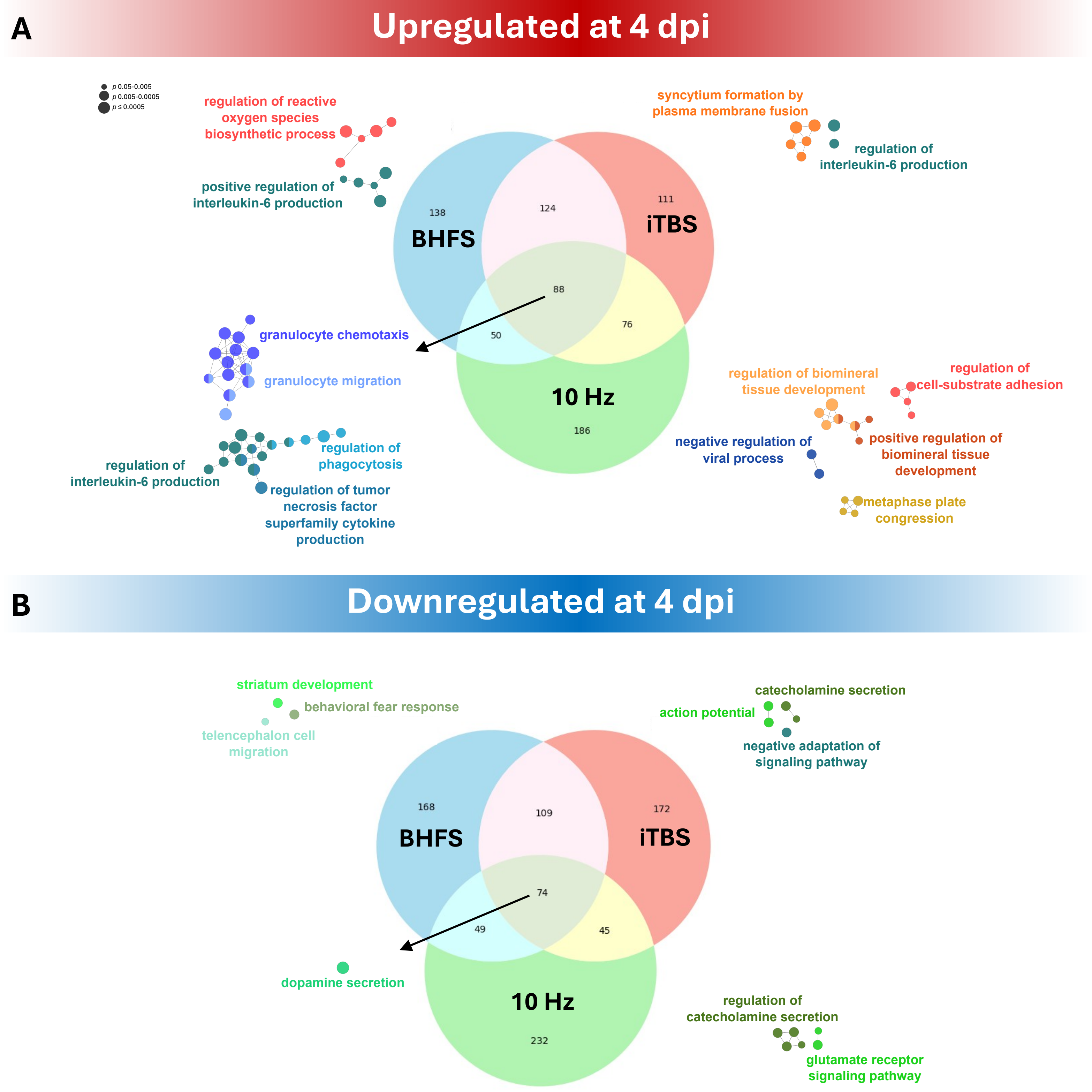

### Supplementary S8

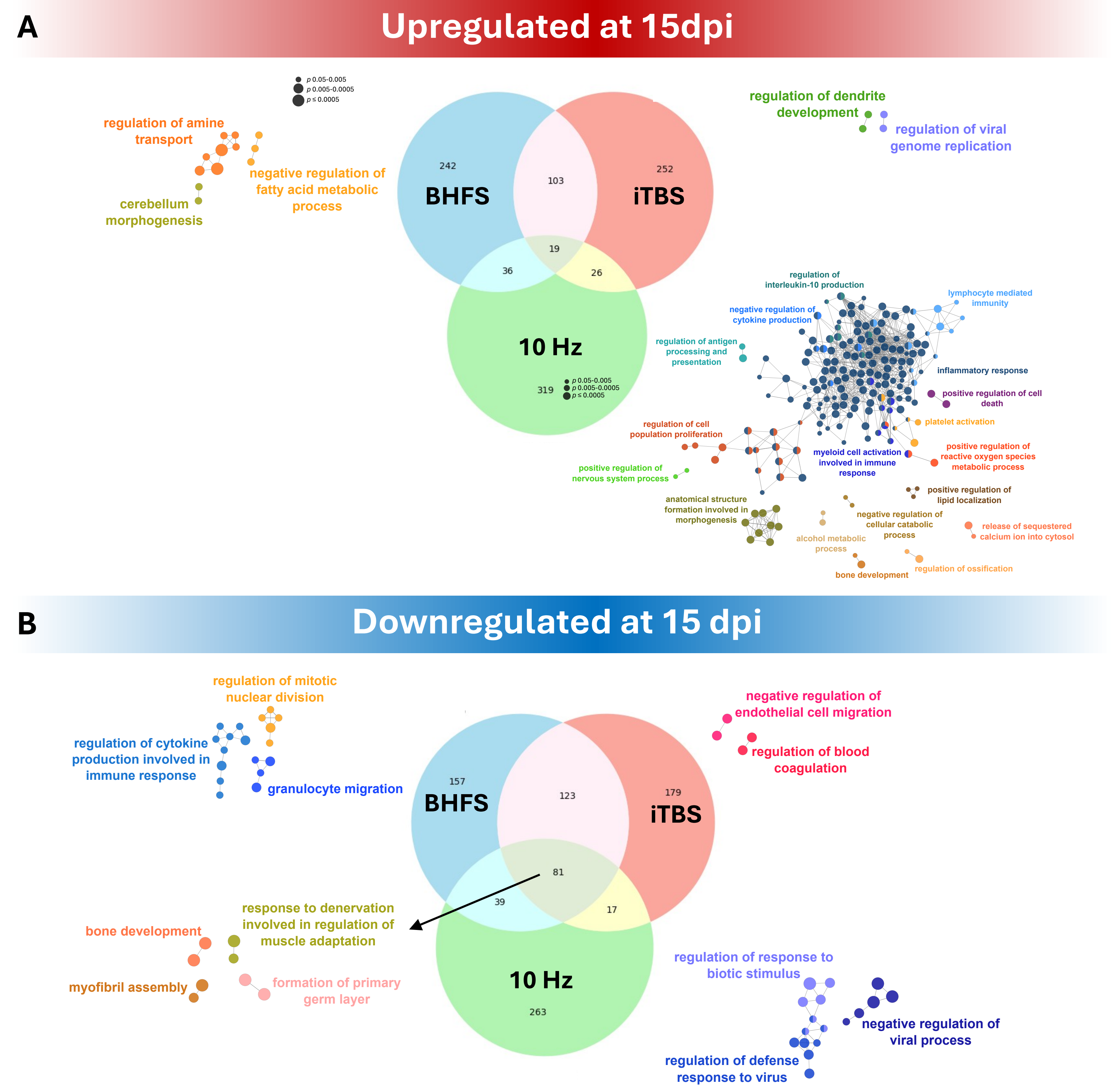

### Supplementary S9

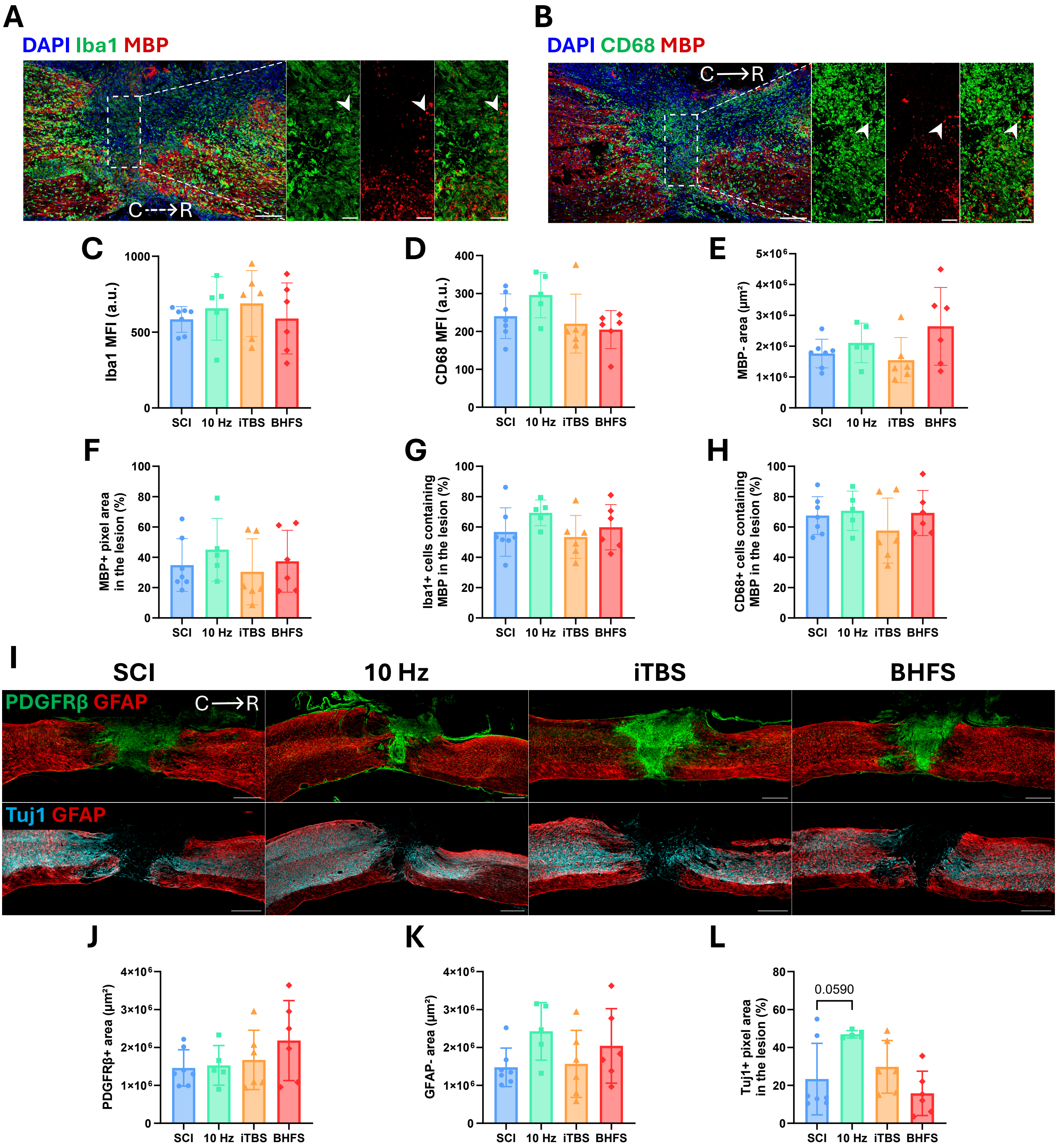

### Supplementary S10

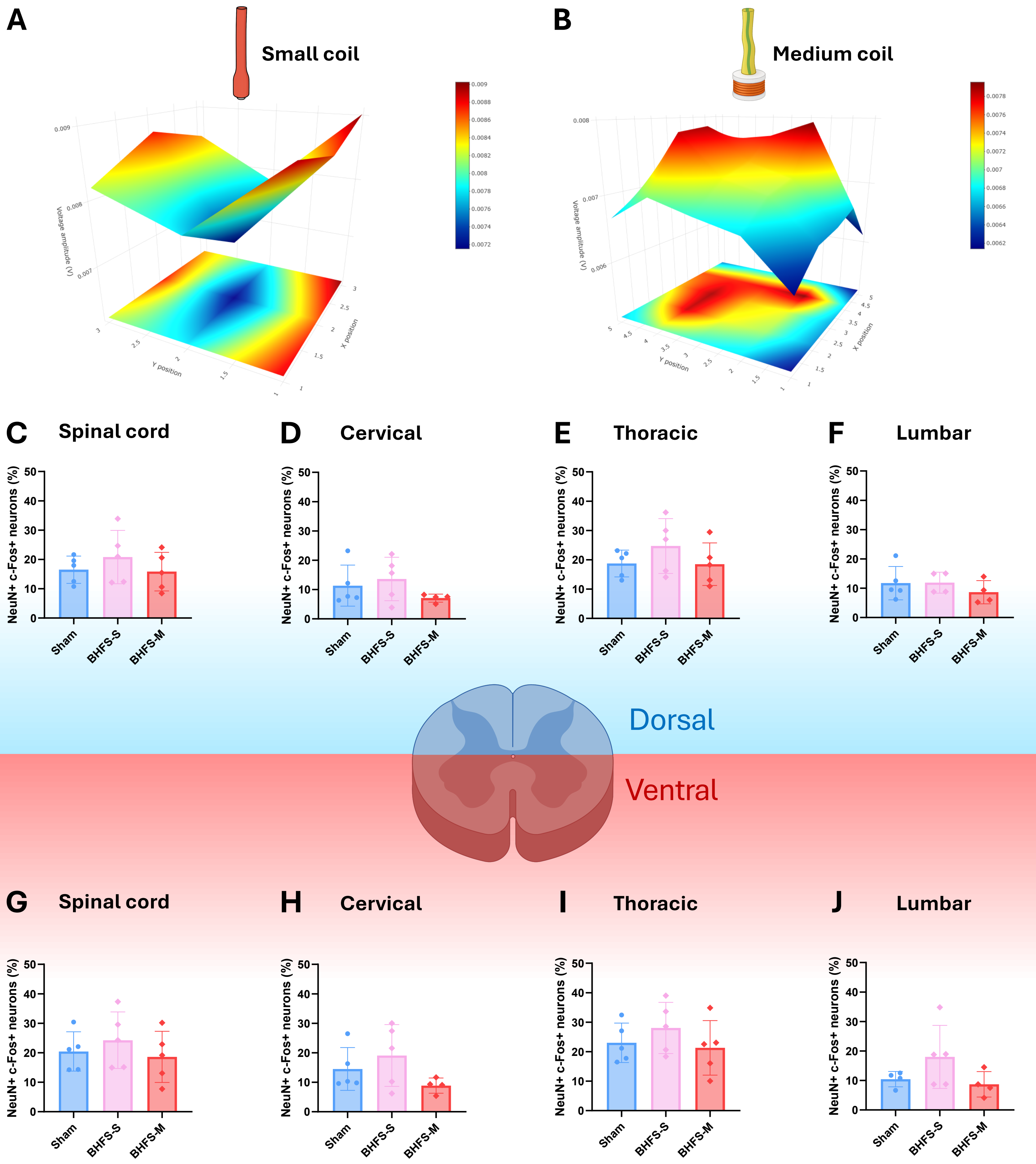
